# The unexpected dual role of S100A9 amyloid protein on neurodegeneration in progressive multiple sclerosis motor cortex

**DOI:** 10.1101/2025.11.21.689707

**Authors:** Jonathan Pansieri, Jonathan Atwood, Marco Pisa, Richard Yates, Gabriele C. DeLuca

**Author notes:** Correspondence to: Jonathan Pansieri, PhD, Principal investigator, UK-MS Early Career Fellow,; Gabriele C. De Luca, MD, DPhil, FRCPath, FAAN, Professor of Clinical Neurology, Nuffield Department of Clinical Neurosciences, University of Oxford. Director, Clinical Neurosciences Undergraduate Education, Oxford Medical School.

## Abstract

Motor cortical inflammation and neurodegeneration are key features of progressive multiple sclerosis (MS), contributing to irreversible motor disability. However, the precise mechanisms leading to neuronal loss remain poorly understood. While chronic inflammation is a hallmark of MS and contributes to disease progression, the factors linking inflammation to neuronal loss in the cortex are not well defined. One candidate is the calcium-binding protein S100A9, a damage-associated molecular pattern (DAMP) protein, thought to be released by stressed cells and infiltrating monocytes. Through its ability to modulate immune responses and influence neuronal survival, S100A9 may sustain chronic inflammation and participate in neurodegenerative processes. Although its role has been explored in other neurological disorders, its contribution to progressive MS remains largely uncharacterised.

Here, we investigated S100A9 expression in the motor cortex of a large post-mortem cohort of MS cases (n=67) and controls (n=9), focusing on its cellular localisation and its relationship with neuronal density and vascular pathology.

S100A9 expression was increased in progressive MS cases compared to controls, predominantly localised to intravascular monocytes and as amyloid-like extracellular plaques surrounding blood vessels. These patterns were associated with neurodegeneration and blood–brain barrier (BBB) disruption, as evidenced by correlations with reduced brain weight, decreased neuronal density, and increased fibrin(ogen) deposition. In contrast, S100A9 was also expressed in microglia, where it correlated with increased neuronal density and reduced fibrin(ogen) deposition. Notably, the phosphorylated form of S100A9, linked to pro-inflammatory signaling, was reduced in microglia but enriched in perivascular regions.

These findings reveal a dual, compartment-specific role for S100A9 in progressive MS, whereby extracellular aggregates may drive neurotoxicity, while microglial S100A9 may confer neuroprotection. Therapeutic strategies targeting extracellular S100A9 while preserving its intracellular functions may offer new opportunities for treating progressive MS.

## Introduction

Multiple sclerosis (MS) is a chronic immune-mediated neurological disorder of the central nervous system characterised by demyelination, inflammation and progressive neurodegeneration, ultimately leading to irreversible disability^1^. In progressive MS, motor cortical damage is a key pathological feature contributing to motor disability^2–4^. However, the mechanisms driving cortical neurodegeneration remain incompletely understood^5^. While cortical demyelination has historically received significant attention, emerging evidence suggests that inflammatory and neurodegenerative mechanisms in the grey matter of the MS motor cortex^6^ can occur independently of demyelination. This has prompted a shift in focus toward additional contributors such as vascular dysfunction, innate immune activation and adaptive immune responses^7,8^. Notably, meningeal inflammation, whether organised into tertiary lymphoid structures or diffuse, has been strongly associated to cortical demyelination and neurodegeneration in progressive MS^9,10^.

Among the key molecular mediators of chronic inflammation are damage-associated molecular patterns (DAMPs) proteins, a group of endogenous molecules released by stressed or damaged cells that activate and sustain immune responses, particularly in relapsing inflammatory disorders^11,12^. One such DAMP, the calcium-binding protein S100A9, has been implicated in several neurodegenerative diseases, including Alzheimer’s (AD)^13^ and traumatic brain injury (TBI)^14^, where it promotes inflammation and amyloid aggregation. Interestingly, S100A9 may also exert protective effects. Indeed, a range of in vitro studies suggest that S100A9 can limit excessive inflammation and tissue damage^15^, promote glial phagocytosis^16^, and inhibit pro-inflammatory macrophage activation^17^. These findings support a context-dependent role for S100A9.

Despite its recognised role in other neurological conditions, the contribution of S100A9 to MS, particularly in its progressive form, remains poorly understood. In animal models of MS (experimental autoimmune encephalomyelitis, EAE), S100A9 contributes to immune cell activation^18^ and active demyelination^19–21^. Moreover, elevated S100A9 levels have been reported in the serum of MS patients during active disease or relapse^22^, suggesting it may be involved in systemic immune activation and CNS infiltration. Together, these findings support a potential role for S100A9 in MS during progression.

In this study, we investigated the expression of S100A9 in the motor cortex of a post-mortem cohort of progressive MS cases (n=67) and non-neurological controls (n=9). We focused on its cellular localisation and its relationship with neuronal density and vascular dysfunction, using fibrin(ogen) accumulation as a surrogate marker of BBB damage in a subset of the cohort (MS, n = 37; control, n = 7). Our findings reveal a dual, compartment-specific role for S100A9 in progressive MS, where extracellular amyloid-like aggregates of S100A9 are associated with neurodegeneration and BBB damage, while microglial expression of S100A9 is linked to preserved neuronal integrity. These results highlight the potential of S100A9 as both a pathological marker and therapeutic target in progressive MS.

## Material and Methods

### Study Population

A post-mortem cohort of pathologically confirmed MS cases (n=67) and non-neurological controls (n=9) from the UK MS Tissue Bank was studied (Table 1). Autopsy material was obtained with the relevant ethics committee approval (REC 08/MRE09/31+5) according to the UK Human Tissue Act (2004). For each case, brain weight was recorded before sampling.

**Table 1.** Demographics of MS and control cohorts. (MS = multiple sclerosis; n/a = not applicable; M = male; F = female; PPMS = primary progressive MS; SPMS = secondary progressive MS; RRMS = relapsing-remitting MS; PM = post-mortem :; *p<0.05; **p<0.01).

|  | Control cases (n=9) | MS cases (n=67) | p-values |
| --- | --- | --- | --- |
| Age at death (y) | 77.5 (range 68-91) | 63.4 (range 40-92) | 0.0014 (**) |
| Duration of disease (y) | N/A | 29.7 (range 11-58) | N/A |
| disease course | N/A | RRMS=4 ; PPMS=7 ;<br>SPMS=50 ; unknown=6 | N/A |
| time onset-to-wheelchair (y) | N/A | 18.9 (range 3-50) | N/A |
| Sex | M = 5 ; F = 4 | M = 17 ; F = 50 | 0.11 |
| Brain weight (g) | 1336 (range 1072-1628) | 1152 (range 800-1557) | 0.0065 (**) |
| PM interval (h) | 26.8 (range 10-52) | 17.0 (range 6-38) | 0.02 (*) |
| **Control and MS sub-cohort used for other markers than S100A9, pS100A9 and TREM2** |  |  |  |
|  | Control cases (n=7) | MS cases (n=37) | p-values |
| Age at death (y) | 78.1 (range 68-91) | 63.3 (range 42-92) | 0.0043(**) |
| Duration of disease (y) | N/A | 29.3 (range 12-56) | N/A |
| disease course | N/A | RRMS=1 ; PPMS=4 ;<br>SPMS=29 ; unknown=3 | N/A |
| time onset-to-wheelchair (y) | N/A | 18.1 (range 4-45) | N/A |
| Sex | M = 4 ; F = 3 | M = 11 ; F = 26 | 0.21 |
| Brain weight (g) | 1320 (range 1072-1628) | 1147 (range 894-1380) | 0.04 (*) |
| PM interval (h) | 28.4 (range 10-52) | 18.2 (range 7-38) | 0.08 |
| **MS sub-cohort at the extremes of microglial S100A9 expression (pS100A9, TREM2 staining)** |  |  |  |
|  | S100A9- microglia (n=10) | S100A9+ microglia (n=10) | p-values |
| Age at death (y) | 61.3 (range 40-82) | 66.6 (range 48-92) | 0.36 |
| Duration of disease (y) | 27.9 (range 8-47) | 32.1 (range 13-58) | 0.42 |
| disease course | PPMS=2 ; SPMS=7 ;<br>unknown=1 | PPMS=1 ; SPMS=8 ;<br>unknown=1 | N/A |
| time onset-to-wheelchair (y) | 18.4 (range 2-39) | 21.0 (range 3-50) | 0.34 |
| Sex | M = 2 ; F = 8 | M = 0 ; F = 10 | 0.47 |
| Brain weight (g) | 1154 (range 885-1557) | 1132 (range 800-1393) | 0.65 |
| PM interval (h) | 15.9 (range 7-27) | 18.0 (range 6-31) | 0.15 |

### Immunohistochemistry and Immunofluorescence

Adjacent sections (6 µm thick) of formalin-fixed paraffin-embedded (FFPE) motor cortical tissues were deparaffinised and conventional antigen retrieval procedures were applied (Supplementary Table 1). FFPE sections were labeled with primary antibodies for S100A9, Neurons (NeuN), fibrin(ogen), microglia (TREM2, Iba1, TMEM119), astrocytes (GFAP), endothelium (CD31/CD34), monocytes (CD163) and phosphorylated S10019 (termed pS100A9), revealed by DAB staining after secondary antibody incubation labelled with horseradish peroxidase (Dako REAL EnVision Detection System, #K5007), and counterstained using haematoxylin, as previously described.^23^ To minimise non-specific background staining, a commercial protein blocker (Protein Bloc Serum-Free, Dako #X0909) was used prior to application of the primary antibody. The omission of primary and secondary antibodies separately served as negative controls.

To investigate the expression of S100A9 in brain cells, fluorescent co-labelling using the same primary antibodies and additional relevant markers as listed in Supplementary Table 1 was performed in a subset of MS cases (n = 2) using relevant secondary antibodies (i.e. Alexa Fluor 488 (anti-rabbit, Invitrogen #A32790), Alexa Fluor 594 (anti-mouse, Invitrogen, # A32744) and Alexa Fluor 647 (anti-chicken, Invitrogen, # A21449), respectively). To minimise autofluorescence, a commercial quencher designed specifically against intrinsic fluorescence of lipofuscin was used (TrueBlack Lipofuscin Autofluorescence Quencher, Biotium #23007).

For dual chromogenic/IF labelling we used for fibrinogen/S100A9 and pS100A9/S100A9 co-labelling, IF staining was performed first for S100A9, followed by antibody stripping using a validated antibody removal kit following previously published procedures^24^, and then re-stained with DAB-based detection for fibrinogen or pS100A9.

### Assessment of Motor Cortical Demyelination

Motor cortical demyelination was defined by complete loss of myelin using PLP immunostaining, as previously published^2,25^. According to well-established classification criteria^26,27^, demyelinated lesions were subsequently classified into type I (leukocortical), type II (intracortical), type III (subpial, layers I-III) and type IV lesions (pancortical, layers I-VI), as illustrated in Supplementary Figure 1.

### Investigation of Motor Cortical S100A9 Expression and Relationships with MS pathology

#### Quantitation of S100A9 expression

##### TOTAL S100A9 EXPRESSION

Total S100A9 expression was quantified in the parenchyma of MS cases and controls within pre-defined spaced trajectories perpendicular to the subpial surface of the motor cortex, as previously described^2^. For each trajectory, 2 fields of view (FOVs) of the same size were applied to motor cortical grey matter (GM) layers, except for layer III, in which 4 FOVs were used due to the larger size of this layer. Cortical layers I-III were designated as the supragranular layers, while layers IV-VI were designated as the infragranular layers. An additional 4 FOVs following the same trajectories were used to quantify the total expression of S100A9 in the white matter (WM) at the border between GM and WM, when applicable. FOVs were defined in non-lesional grey matter (NLGM) and white matter (NLWM). Of note, 7 cases from the MS cohort show total subpial demyelination, meaning that layer I-III assessment for NLGM in these layers was not possible in this subset of cases.

For GM lesions, FOVs were defined using the same strategy as for NLGM and compared to corresponding NLGM cortical layers in the immediate border-lesional area – from lesion edge up to 300 µm and NLGM –300 to 600 µm from lesion edge. For WM lesions, 5 FOVs per lesion were arranged around the centre of the lesion and also compared to the immediate border-lesional area – from lesion edge up to 300 µm and NLGM –300 to 600 µm from lesion edge. In demyelinated cases, 78 lesions were identified, wherein 43 lesions qualified given availability of lesional, border-lesional and NLGM areas.

Employing this standardized approach, more than 14,000 FOVs were analyzed to investigate S100A9 expression. Each case was scanned using Aperio ScanScope AT Turbo digital scanner at 40X magnification.

Total S100A9 expression was quantified in each FOV by an established semi-automatic color-based extraction method using QuPath, and expressed as area coverage (%), i.e. chromogen-positive area divided by FOV area.

S100A9 was predominantly found in intravascular monocytes, extracellularly deposited around the vessels, and in microglia, which prompted additional quantitative and semi-quantitative assessments for each compartment in both MS cases and controls. For each assessment, data was averaged in supragranular layers (layers I-III) and infragranular layers (layer IV-VI).

##### QUANTITATION OF S100A9+ MONOCYTE-ASSOCIATED VESSELS

Vessels in the longitudinal plane containing S100A9+ monocytes with an elongated sausage-shaped morphology (as previously described^20^) were manually counted in each NLGM FOV (Supplementary Figure 2) and expressed as vessels/mm^2^. Vessels containing more than one S100A9+ monocyte were counted as a single positive vessel. Using this method, more than 5,000 S100A9+ monocyte-associated vessels were counted.

##### QUANTITATION OF PERIVASCULAR S100A9 PLAQUES

Immunopositive perivascular deposits of S100A9 around blood vessels were defined as perivascular S100A9 plaques and counted in both longitudinal and transverse blood vessels (Supplementary Figure 3) in each NLGM FOV. The distribution of perivascular S100A9 plaques were measured within a predefined 70 µm radial distance from the vessel wall, standardised to ensure consistency, and expressed as plaques/mm^2^. Only plaques with a diameter larger than 5 µm were included to ensure consistency and relevance in the measurement. Using this method, more than 1,000 plaques were counted.

##### SEMI-QUANTITATIVE MICROGLIAL SCORING OF S100A9

In each FOV, a semiquantitative scale between 0 and 2 was used to assess S100A9 expression in microglial compartments. Scoring was defined as 0 (no staining), 1 (sparse and predominantly vessel-associated staining of microglial processes) and 2 (frequent perivascular and parenchymal expression in microglial soma and processes).

Scoring was validated for reliability within rater (test-retest) and between two independent raters on 1,421 FOVs from 10 MS cases using Cohen’s kappa inter-rater agreement test (both reliability tests, κ(glial) >0.80).

#### Quantitation of neuronal density and fibrin(ogen) accumulation

The relationship between S100A9 and MS motor cortical pathology was evaluated in a subset of cases (37 MS cases, 7 controls) that were previously characterised for various markers of interest^2,25,28^.

##### NEURONAL DENSITY

Neuronal density was quantified in each FOV by manually counting the number of NeuN+ cells in layers III and V, which were chromogen-positive, expressed as cells/mm^2^, as previously described^23^. Using this method, more than 50,000 NeuN+ neurons were counted.

##### FIBRIN(OGEN) DEPOSITION

Fibrin(ogen) deposition was quantified in each FOV by an established semi-automatic color-based extraction method using QuPath software, and expressed as chromogen-positive pixels/mm^2^, as previously described^2^.

#### Quantitation of pS100A9 and TREM2

##### PHOSPHORYLATED S100A9 AND TREM2 EXPRESSION

Motor cortical pS100A9 and TREM2 expression was assessed in adjacent sections of a subset of MS cases at the extremes of S100A9 microglial expression (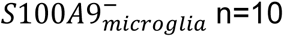 average score 0; 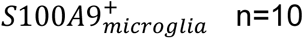, average score 1.2). FOVs were examined along the same trajectories as for S100A9 expression, using the Interactive Image Alignment tool in QuPath. pS100A9 and TREM2 expression were assessed using the same semiautomatic color-based extraction method described above for S100A9, expressed as % area coverage.

### Statistical analyses

Clinical features comparing multiple sclerosis (MS) and control groups were assessed using the non-parametric Mann-Whitney U test.

A Generalized Linear Model (GLM) was conducted to evaluate the impact of disease status (MS vs. control) on S100A9 expression in NLWM and NLGM, including post-mortem interval (PMI) and age-at-death as covariates. Despite the inclusion of these covariates, neither PMI nor age-at-death significantly influenced the S100A9 expression results.

The S100A9 dataset was log-transformed to address non-normality prior to conducting a stepwise regression analysis with age, post-mortem interval, and sex as predictors. The stepwise regression analysis indicated that none of these predictors significantly impacted S100A9 expression, leading to their exclusion from further analysis.

To compare S100A9 scores and densities, fibrinogen expression, and neuronal density between MS and control groups, the Mann-Whitney U test was employed. Analyses across cortical layers involved multiple comparisons, where the Bonferroni-Dunn correction was applied to adjust for Type I errors.

Paired data for non-lesional vs. lesional areas were analysed using one-way ANOVA with Geisser-Greenhouse correction and Bonferroni’s multiple comparisons test; when relevant. Correlation analyses to evaluate relationships between S100A9 expression and clinical features, fibrinogen expression, and neuronal density were performed using Spearman rank-correlation coefficients.

Data are presented with ± mean deviation of the mean. Two-tailed tests were applied where appropriate, with p-values < 0.05 considered statistically significant. Statistical analyses were conducted using GraphPad Prism (version 9.5.1) and SPSS (version 29).

## Results

### Demographic features of MS cohort

Clinical details of the cohort can be found in Table 1. Most MS cases were female (female, 50 of 67 = 74.6%, male, 17 of 67 = 25.4%) and the median age was 63.4 years. Most MS cases were classified as secondary progressive (74.6%) or primary progressive (10.4%), with a small number of relapsing-remitting (6.0%) and unknown clinical course (9.0%). Brain weight was reduced in MS compared with controls (in grams; MS: 1145, CI95% 1110-1181; control: 1336, CI95% 1189-1484; p=0.0065). MS cases died younger than controls (in years; MS: 63.6, CI95% 60.6-66.5 versus control: 77.6, CI95% 72.0-82.9; p=0.0014), and post-mortem interval was shorter in MS cases compared to controls (in hours; MS: 16.8, CI95% 15.1-18.5 versus control: 26.8, CI95% 15.7-37.9; p=0.02). No difference in sex was found between MS cases and controls.

Demyelinated lesions were detected in 64.2% of MS cases (43 out of 67). In particular, type I lesions were found in 1.5%, type II in 3.0%, type III in 62.6%, and type IV in 10.5% of MS cases. WM lesions were detected in 7.5% of MS cases. No demyelinated lesions were found in controls.

### S100A9 is expressed in various cell types and is extracellularly deposited into plaques in MS motor cortex

We performed conventional immunohistochemistry and double-labelling immunofluorescence to identify the specific cell types which express S100A9 in the motor cortex of MS cases and controls (Figure 1A-G, Supplementary Figure 4).

**Figure 1.**
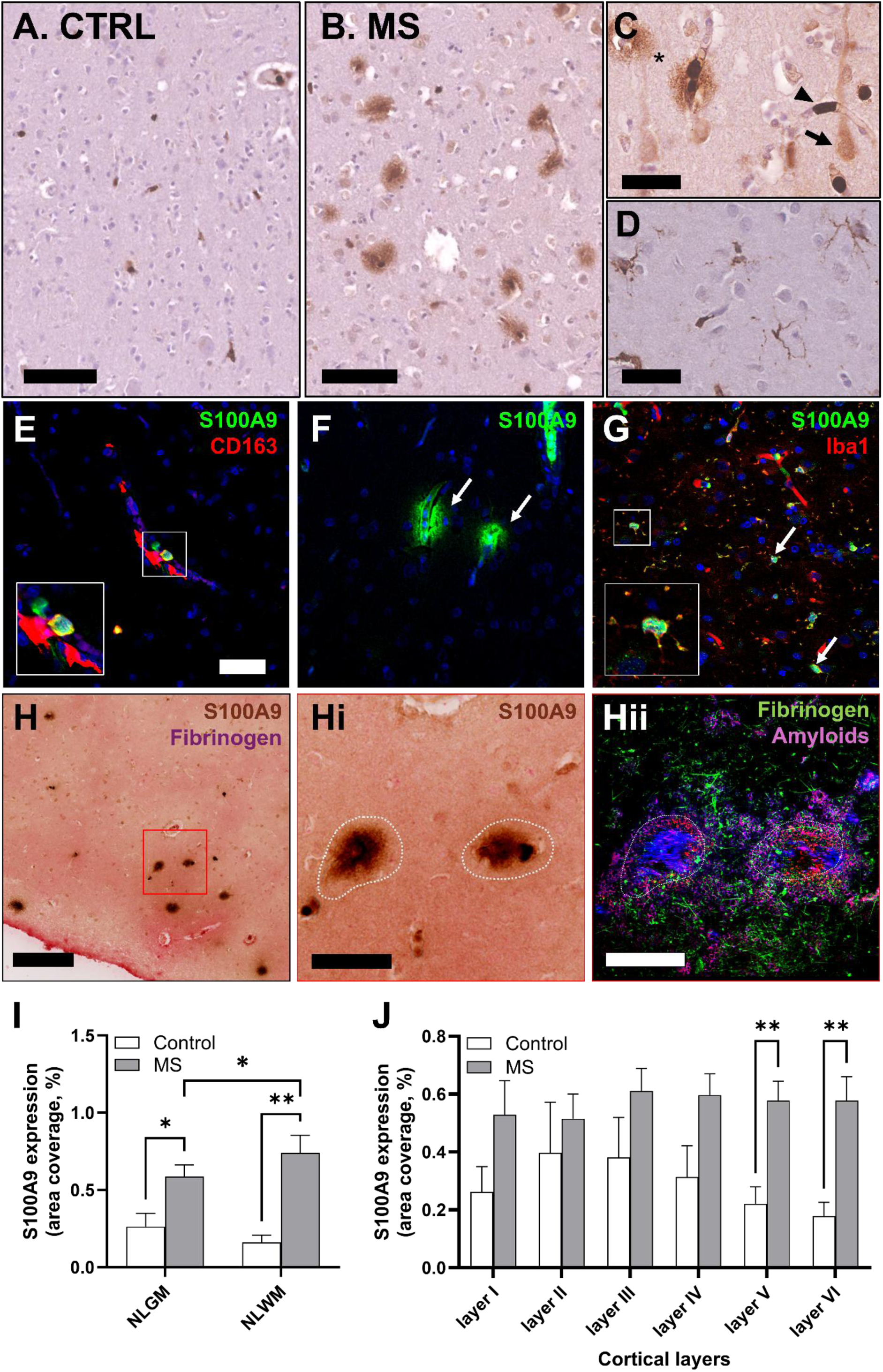
S100A9 expression in MS and control motor cortex. **(A-B)** Representative S100A9 expression in **(A)** control and **(B)** MS non-lesional grey matter of the motor cortex. **(C)** In both MS and controls, S100A9 was found in vessels (arrowhead). In MS, S100A9 was also extracellularly deposited around vessels (arrow) sparsely expressed in neurons (asterisk), and **(D)** expressed in microglia. **(E-G)** For each pattern, fluorescent labelling identified S100A9 (green) to be expressed **(E)** in intravascular monocytes (CD163, red), **(F)** perivascularly deposited (arrows) and **(G)** expressed in microglia/macrophages (Iba1, red, arrows). **(H, Hi)** Extracellular S100A9 (brown) accumulates around fibrinogen (purple, green), a surrogate of blood-brain barrier damage. Importantly, S100A9 deposits show **(Hii)** typical amyloid-related fluorescence properties (blue, λ_ex_=380nm, λ_em_=440nm); red, λ_ex_ =640nm, λ_em_ =690nm). (Scale bars −500µm). **(H)** S100A9 was increased in NLWM and NLGM of MS cases compared with controls. S100A9 was also increased in NLWM compared to NLGM, in MS cases only. **(I)** The increase of S100A9 expression in NLGM was restricted to cortical layers V and VI. (Results are presented as mean ± SEM; *p<0.05; **p<0.01; **** p<0.0001; MS = multiple sclerosis; CTRL = control; NLGM = non-lesional grey matter, NLWM = non-lesional white matter; scale bars in **(A, B)** are 100 µm, in **(C, D)** – 40µm, in **(E-G, Hi, Hii)** – 50 µm and **(H)** – 500 µm; 400X magnification; each **(E-G)** fluorescent picture is supplemented by Hoechst staining in blue, which stain cell nuclei.)

In controls, the predominant S100A9 expression pattern comprised intravascular cells with an elongated sausage-shaped morphology (Figure 1A), showing cytoplasmic S100A9 immunoreactivity surrounded by endothelial CD31/CD34+ cells (Supplementary Figure 4). This pattern remained in most MS cases, supplemented by extracellular, perivascular S100A9 deposition in the form of dense, plaque-like deposits or diffusely expressed, adjacent to small vessels in the extracellular compartment (Figure 1B), with a diameter of approximately 20–70μm. In MS and controls, we also observed sparse neuronal expression of S100A9, exhibiting predominant cytoplasmic expression in the form of granules, and co-locating with NeuN neuronal marker (Figure 1C, Supplementary figure 4). Another distinct S100A9 pattern was a glial-like morphology of S100A9 cells within the parenchyma, characterised by ramified processes and moderate S100A9 immunoreactivity in the soma and proximal processes (Figure 1D).

Using immunofluorescence, intravascular S100A9+ sausage-shaped cells were identified as monocytes by co-localisation with CD163+ monocytes (Figure 1E). Perivascular S100A9 plaques was also confirmed by fluorescence imaging (Figure 1F), and glial-like parenchymal S100A9 expression was identified as Iba1+ and TMEM119+ microglia/macrophages (Figure 1G, Supplementary Figure 4). Of note, S100A9 was almost absent in astrocytes (Supplementary Figure 4).

Importantly, we also confirmed the S100A9 deposition around areas of vascular dysfunction using fibri(nogen), a surrogate of blood-brain barrier damage (Figure 1H), showing the amyloid nature of S100A9 by specific fluorescence signatures we previously described^29^ and allowing us to define them as plaques.

Of note, in the white matter of both MS and control cases, S100A9 was additionally observed in oligodendrocyte-like cells displaying a characteristic “fried-egg” morphology, with a small, round, and densely stained nucleus surrounded by a clear perinuclear halo and thin cytoplasmic rim (data not shown).

### S100A9 is upregulated in MS motor cortex compared to controls

Quantitative measures of S100A9 expression were performed in MS cases and controls. A summary of results for each cortical layer is available in Supplementary Table 2. Importantly, stepwise regression analysis showed no impact of sex, age-at-death and post-mortem interval in our analyses, as outlined in the methods.

#### EXTENT OF S100A9 EXPRESSION

The extent of total S100A9 expression in non-lesional grey matter (NLGM), non-lesional white matter (NLWM) and demyelinated areas was quantified in MS cases and controls (Figure 1H-I, Supplementary Figure 5).

Relative to controls, S100A9 expression was increased in both NLWM and NLGM of MS motor cortex (Figure 1I; area coverage in %; NLWM, MS: 0.74, CI95% 0.51 – 0.97 vs controls: 0.16, CI95% 0.05 – 0.27; Waldχ^2^ 7.9, p=0.002; NLGM, MS: 0.59, CI95% 0.44 – 0.74 vs controls: 0.22, CI95% 0.06 – 0.38; Waldχ^2^ 5.05, p=0.016). Layer-by-layer quantification showed that total S100A9 expression was more abundant specifically in infragranular layers (Figure 1J, MS Vs CTRL, layer I, p=0.38; layer II, p=0.33; layer III, p=0.12; layer IV, p=0.08; layer V, p=0.007; layer VI, p=0.009). Removal of MS cases, who died by sepsis, did not impact our findings, despite the well-known contribution of S100A9 in this condition^30–32^.

In MS cases, total S100A9 expression was increased in NLWM compared to NLGM (Area coverage, %; MS: NLWM Vs NLGM: p=0.02; Control: NLWM Vs NLGM: p=0.15).

In MS, no difference in total S100A9 expression was found between non-lesional, border-lesional and lesional areas, taking into account all lesion types and each type of lesion separately (Supplementary Figure 5; all comparisons, p>0.4).

No relationship was found between total S100A9 expression and post-mortem interval, brain weight, age at death, disease duration, or time from onset to wheelchair in MS or controls, where applicable.

#### S100A9 PREDOMINANT PATTERNS

Although S100A9 expression in intravascular monocytes was consistently found in NLGM of both MS and controls, microglial S100A9 and perivascular S100A9 patterns were reciprocally expressed in our cohort (Figure 2). Thus, when S100A9 was found in microglia, perivascular S100A9 plaques were almost absent and vice versa, supplemented by cases showing almost only intravascular S100A9 expression. This clear separation enabled a qualitative classification of cases based on dominant S100A9 expression profiles, namely, intravascular, perivascular, or microglial S100A9 patterns.

**Figure 2.**
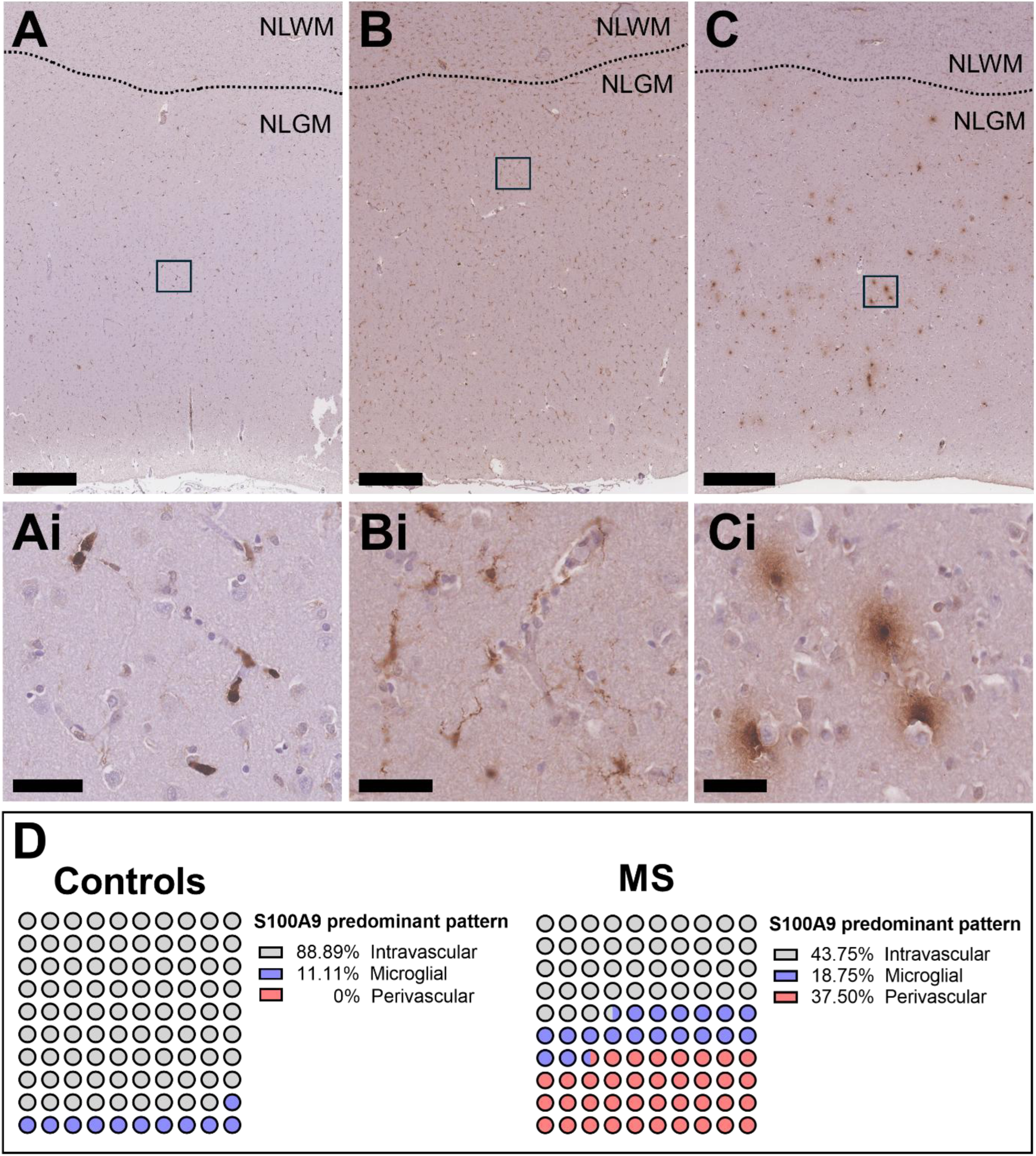
Heterogeneity of S100A9 expression in MS cases and controls motor cortex. Despite a consistent expression in intravascular monocytes throughout our cohort, we observed almost exclusive case-dependent patterns in NLGM. Some cases show **(A)** predominant S100A9+ intravascular monocytes, **(B)** predominant microglial S100A9 expression and **(C)** predominant perivascular S100A9 plaques. Each pattern is accompanied by magnified pictures in **(Ai), (Bi) and (Ci),** respectively. **(D)** Proportion of controls (n=9) and MS cases (n=64) showing each predominant pattern. Of note, two MS cases were excluded from this analysis showing intra-case heterogeneous patterns. (Scale bars are (A-C) 500 µm and (Ai,Bi,Ci) 50µm; NLWM=non-lesional white matter; NLGM = non-lesional grey matter)

Based on this morphological and cellular characterisation, 43.8% of MS cases were intravascular S100A9 predominant (versus 89% in controls), 37.5% were perivascular S100A9 predominant (versus 0% in controls) and 18.8% microglial S100A9 predominant (versus 11.1% in controls).

Of note, the increase in total S100A9 expression observed in NLGM from MS cases compared to controls was driven by MS cases showing case-dependent S100A9 perivascular and S100A9 microglial patterns (Supplementary Figure 6).

### S100A9 expression within and around vessels is increased and relates to reduced brain weight in MS motor cortex

Quantitative and semi-quantitative measures of S100A9 expression were performed focusing on supragranular and infragranular layers, given the increase of total S100A9 expression in infragranular layers in MS cases compared to controls.

#### DENSITY OF S100A9+ MONOCYTE-ASSOCIATED VESSELS

S100A9+ monocyte-associated vessel density was increased in the infragranular cortical layers of MS cases compared to controls (Figure 3A-B; vessels/mm^2^, MS Vs control; supragranular layers: p=0.47; infragranular layers: p=0.009). Importantly, S100A9+ monocyte-associated vessel density was related to reduced brain weight in MS cases, in particular in infragranular cortical layers (Figure 3C, all layers, r=−0.26, p=0.036; supragranular layers, r=−0.24, p=0.07; infragranular layers, r=−0.36, p =0.004).

**Figure 3.**
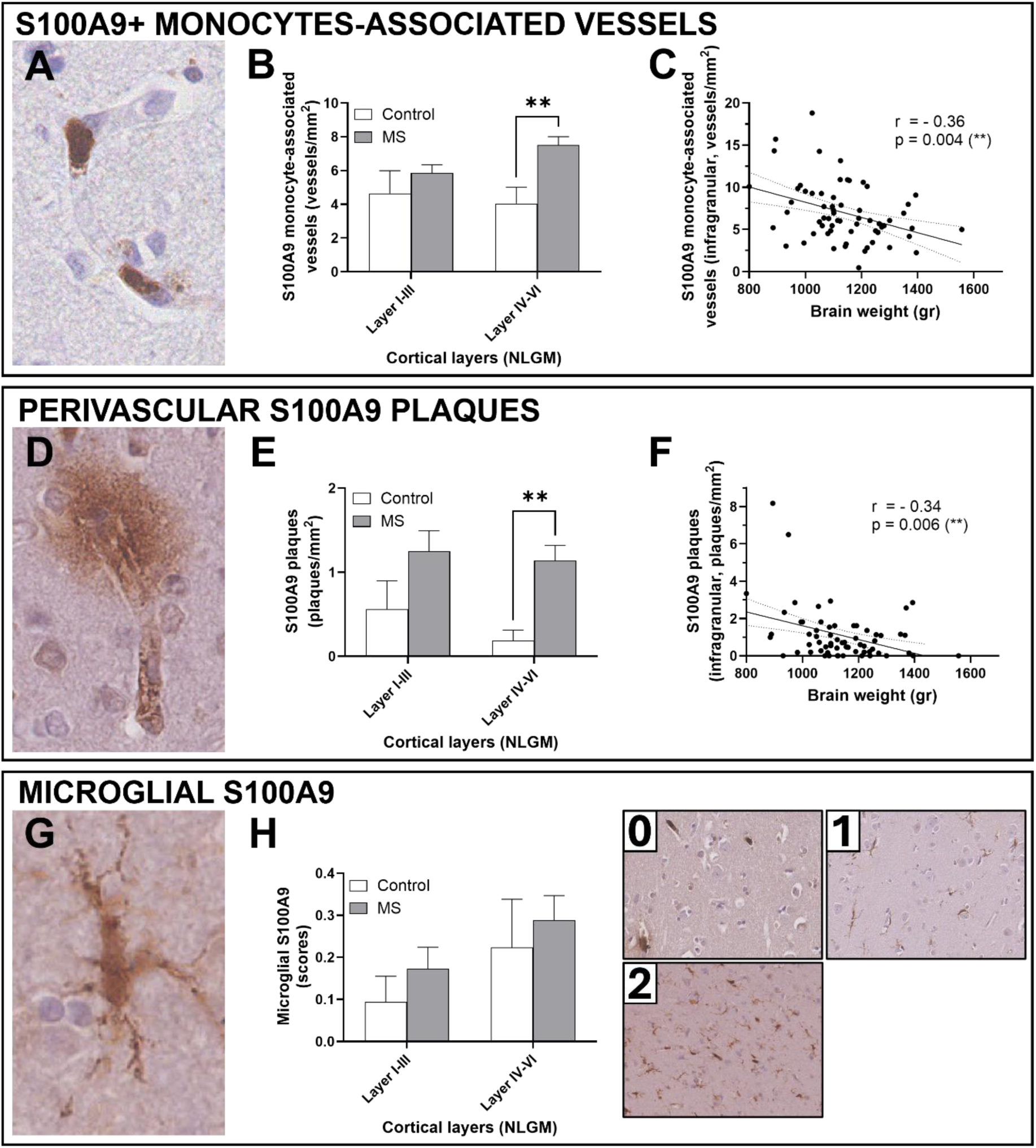
Vascular-related and microglial S100A9 expression in MS and control motor cortex. **(A)** S100A9+ monocyte-associated vessel density **(B)** was increased in MS cases compared to controls, a finding restricted to infragranular layers (layers IV-VI). **(C)** S100A9+ monocyte-associated vessels density in MS infragranular layers was associated with reduced brain weight in MS (right panel). **(D)** Perivascular S100A9 plaque density **(E)** was increased in MS compared to controls, restricted to infragranular layer (layer IV-VI). **(F)** Perivascular S100A9 plaque density in MS infragranular layers was associated with reduced brain weight in MS. **(G)** Microglial S100A9 was **(H,middle panel)** comparable in MS compared to controls across the cortical layers**. (H, right panel)** The scoring for microglial S100A9 is illustrated in left panels and categorised as follows: (0) indicated no staining, (1) represented sparse and predominantly vessel-associated staining of microglial processes, and (2) denoted frequent perivascular and parenchymal expression in microglial soma and processes. (results presented as ± SEM; *p<0.05; **p<0.01; ***p<0.001; gr = grammes; MS = multiple sclerosis; NLGM = non-lesional grey matter).

No relationship was found between S100A9+ monocyte-associated vessel density and other clinical features in MS cases or controls.

Of note, the increase in S100A9+ monocyte-associated vessel density observed between MS cases and controls was conserved throughout the case-dependent patterns observed in MS (Supplementary Figure 6).

#### DENSITY OF PERIVASCULAR S100A9 PLAQUES

Perivascular S100A9 plaque density was greater in MS cases compared to controls, in particular in infragranular layers (Figure 3D-E; plaques/mm^2^, MS vs control; supragranular layers: p=0.14; infragranular layers: p=0.003). Perivascular S100A9 plaque density was related to reduced brain weight in MS cases, a finding restricted to infragranular cortical layers (Figure 3F, all layers, r=−0.26, p=0.036; supragranular layers, r=−0.22, p=0.10; infragranular layers, r=−0.34, p =0.006).

Of note and as expected, the increase in perivascular S100A9 plaques observed between MS cases and controls was driven by MS cases showing case-dependent S100A9 perivascular pattern (Supplementary Figure 6).

#### MICROGLIAL S100A9 SCORING

No overall difference in microglial S100A9 expression was found in MS cases compared to controls (Figure 3G-H, both comparison, p>0.7). However, we found an increase in total S100A9 expression observed between MS subgroups with a microglial S100A9 predominant pattern and controls (Supplementary Figure 6).

Total S100A9 expression, S100A9+ monocyte-associated vessel density, perivascular S100A9 plaque density, and microglial S100A9 did not relate to other clinical features in MS cases or to demographic features in MS cases or controls.

In MS, no difference in all S100A9 measures was found between non-lesional, border-lesional and lesional areas, taking into account all lesion types and each type of lesion separately (data not shown).

#### RELATIONSHIPS BETWEEN S100A9 PATTERNS

Correlation analysis revealed that total S100A9 expression was associated with all other S100A9 measures, including density of monocytes-associated vessels, perivascular plaque density, and microglial scoring. Intravascular S100A9 expression also correlated with perivascular S100A9, suggesting a possible continuum between these vascular compartments. Microglial S100A9 correlated with total and intravascular S100A9, but not with perivascular S100A9 plaque density. This dissociation supports our qualitative observations that microglial and S100A9 plaques are mutually exclusive and may reflect distinct compartmentalisation of S100A9 in MS pathology. (Supplementary Figure 7).

### S100A9 expression relates to neuronal density in MS motor cortex

To further decipher the influence of S100A9 in MS motor cortex pathology, we used a subset of well-characterised MS cases derived from the original cohort^2,25,28^, matched for clinical features (Table 1). Similar levels of S100A9 expression were found in both cohorts (Original MS cohort Vs MS subset: p=0.54) and S100A9 expression followed a similar distribution in both groups (Supplementary Figure 8). Quantitative and semi-quantitative measures of S100A9 were compared with NeuN+ neuronal density in MS cases and controls, with a focus on infragranular layers where S100A9 expression was increased in MS (Figure 4A-C).

**Figure 4.**
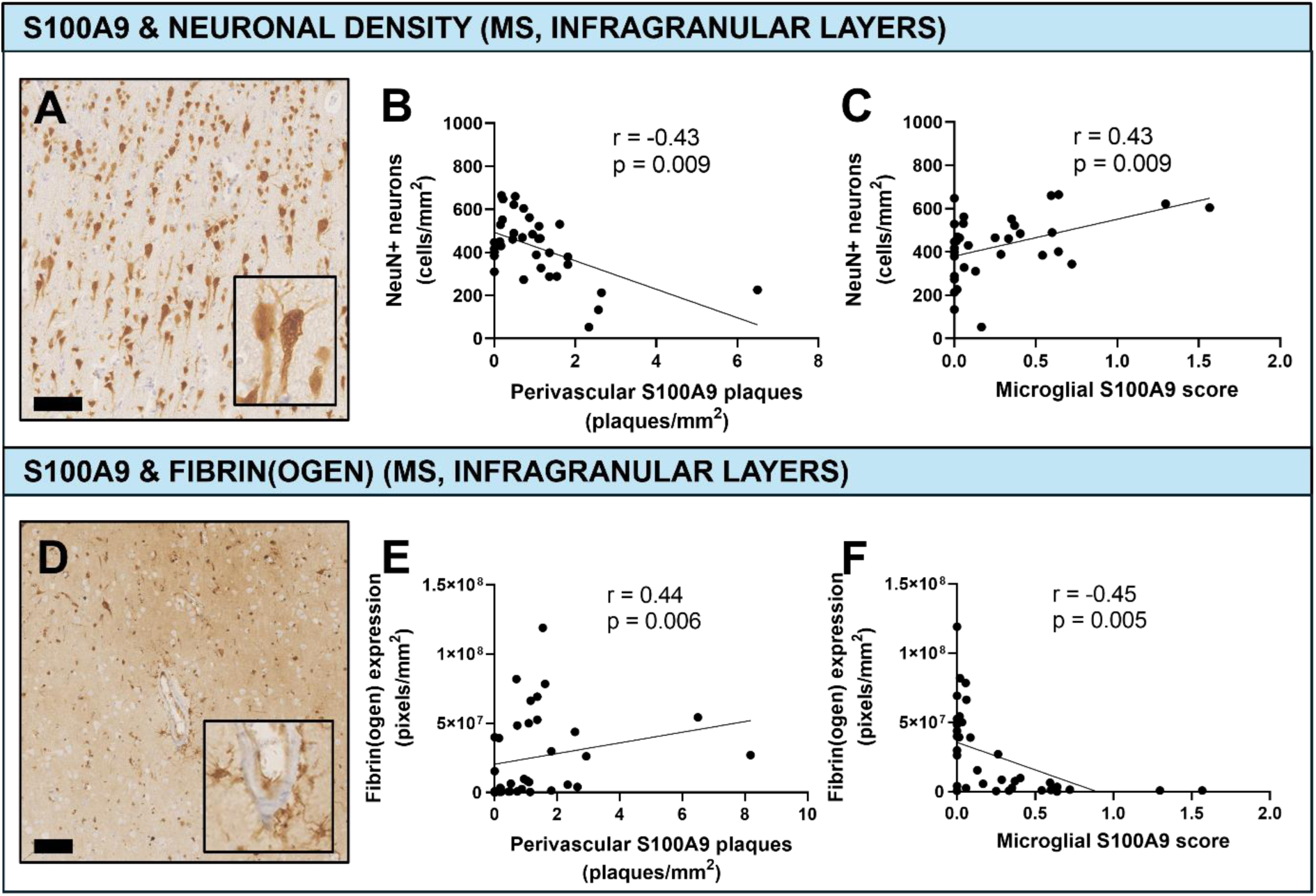
S100A9 expression and MS pathology. **(A)** Representative NeuN+ staining in a MS case. **(B)** Perivascular S100A9 plaques density relates to reduced NeuN+ neuronal density in MS motor cortex infragranular layers. **(C)** Conversely, microglial S100A9 relates to increased NeuN+ neuronal density in MS motor cortex infragranular layers. **(D)** Representative fibrin(ogen) staining in a MS case showing diffuse and astrocytic staining around a vessel. **(E)** Perivascular S100A9 plaques density relates to increased fibrin(ogen) accumulation in MS motor cortex infragranular layers. **(F)** Conversely, microglial S100A9 relates to reduced fibrin(ogen) accumulation in MS motor cortex infragranular layers. (**p < 0.01, scale bars 100µm).

#### NEURONAL DENSITY

No difference was found in NeuN+ neuronal density between MS cases and controls in the NLGM, as previously published^23,25^ (Supplementary Figure 9). However, reduced NeuN+ neuronal density was associated with perivascular S100A9 plaque density, restricted to infragranular layers (Figure 4B, all layers; r=−0.31, p=0.07; supragranular layers, r=−0.23, p=0.21; infragranular layers, r=−0.43, p=0.009). Conversely, increased NeuN+ neuronal density was associated with microglial S100A9, also restricted to infragranular layers (Figure 4C, all layers; r=0.37, p=0.026; supragranular layers, r=0.31, p=0.08; infragranular layers r=0.43, p=0.009). Similar findings were observed by comparing case-dependent S100A9 patterns in MS cases compared to controls (Supplementary Figure 9).

No relationships between neuronal density and total S100A9 expression or S100A9+ monocyte-associated vessel density were found throughout the layers in MS cases (Supplementary Figure 9). No relationship between neuronal density and measures of S100A9 expression was found in controls (all correlations, p>0.30; data not shown).

### S100A9 expression relates to fibrin(ogen) accumulation in MS, a surrogate of BBB damage

Given that the pattern of S100A9 expression in MS cases is consistent with the potential extravasation of S100A9 from blood vessels, S100A9 measures were compared to those of fibrin(ogen), a surrogate of BBB damage, in MS cases and controls (Figure 4D-F, Supplementary Figure 10).

#### FIBRIN(OGEN) DEPOSITION

Fibrin(ogen) deposition was increased in MS compared to controls (Supplementary Figure 10), as previously published^2^. Increased fibrinogen expression was associated with perivascular S100A9 plaque density (Figure 4E, all layers; r=0.46, p=0.004; supragranular layers, r=0.43, p=0.01; infragranular layers, r=0.44, p=0.005). Conversely, reduced fibrinogen expression was associated with microglial S100A9, restricted to infragranular layers (Figure 4F, all layers; r=−0.40, p=0.015; supragranular layers, r=−0.01, p=0.95; infragranular layers, r=−0.45, p=0.005). Similar findings were observed by comparing case-dependent S100A9 patterns in MS cases compared to controls (Supplementary Figure 10), and comparing extracellular deposition of fibrinogen as previously reported and S100A9 measures (data not shown). Of important note, we confirmed co-location between extracellular fibrinogen deposition and perivascular S100A9 plaques (Supplementary Figure 11).

No relationships between fibrinogen expression and total S100A9 expression nor monocyte-associated vessel density were found throughout the cortical layers in MS (Supplementary Figure 10). No relationship between fibrinogen expression and measures of S100A9 expression was found in controls (all correlations, p>0.15; data not shown).

### Microglial S100A9 is predominantly non-phosphorylated and relates to a TREM2+ protective phenotype in MS

Given the unexpected associations between MS pathology and S100A9 expression in microglial compartment, pS100A9 was compared to S100A9 expression in the NLGM of MS cases lying at the extremes of microglial S100A9 scores (lowest extreme: 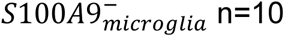; highest extreme: 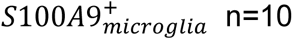; Figure 5, Table 1).

**Figure 5.**
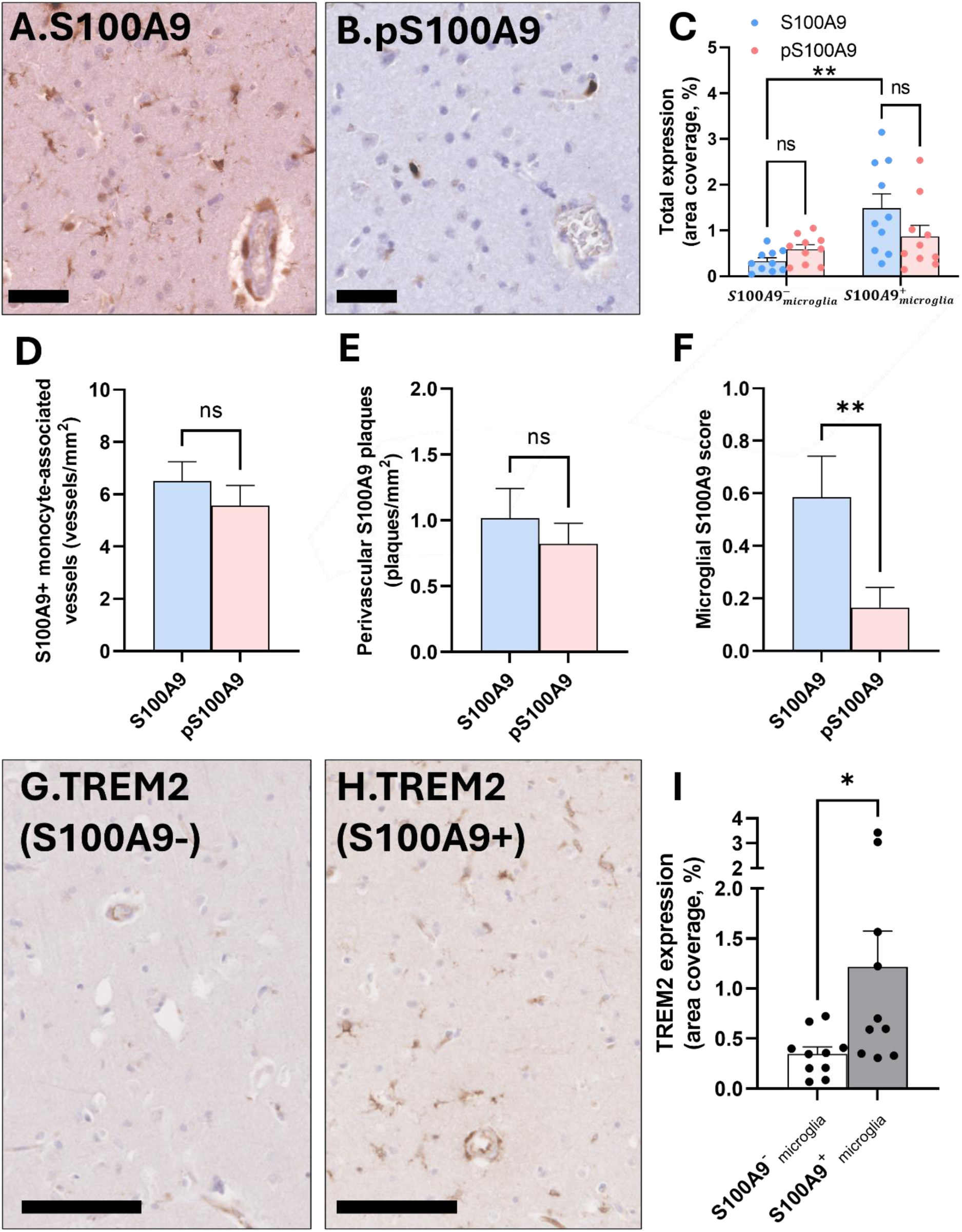
Microglial S100A9 is predominantly non-phosphorylated and associates with TREM2+ microglia in MS motor cortex. **(A-B)** Representative pictures of the same area in adjacent sections of a MS case showing **(A)** S100A9 is expressed in vessels and microglia, while **(B)** pS100A9 is only expressed in vessels. **(C)** In the extremes of microglial S100A9 expression (lowest extreme: 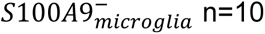; highest extreme: 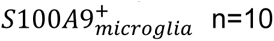, no difference in total S100A9 expression compared to total pS100A9 expression was found. **(D-E)** No difference comparing S100A9+ and pS100A9+ monocyte-associated vessel densities was found (n=20), as well as comparing perivascular S100A9 and pS100A9 plaque densities (n=20). **(F)** However, microglial pS100A9 was specifically reduced compared to microglial S100A9 (n=20, p=0.002). **(G–H)** TREM2+ microglia at the lowest **(G)** and highest (H) extremes of microglial S100A9 expression of our MS cohort. **(I)** Quantification of TREM2 expression at the extremes of microglial S100A9 expression demonstrates an increase in TREM2+ microglia in MS cases with high microglial S100A9 (n=20, p=0.035). (Results presented as ± SEM; **p < 0.01, scale bars 50µm in A-B, 100µm in G-H).

No difference was found between pS100A9 expression and S100A9 expression in MS cases lying at the extremes of microglial S100A9 expression, considering total expression, plaque density nor monocyte-associate vessels density. However, a specific reduction in microglial pS100A9 compared to microglial S100A9 was found (Figure 5A-F, Supplementary Figure 12). Of note, when expressed in glial cells, pS100A9 was mainly observed in perivascular macrophages.

To further characterise microglial S100A9 expression, we assessed TREM2, a marker of phagocytic microglia considered to reflect a protective or reparative phenotype in the extremes of microglial S100A9 expression.

TREM2+ microglia were identified in the MS motor cortex, displaying small, rounded somata with thickened, retracted processes (Figure 5G–H). Quantitative analysis revealed an increase in TREM2+ microglia in MS cases with high microglial S100A9 expression compared to those with low expression (Figure 5I).

## Discussion

This study investigated the expression of S100A9 in the motor cortex of progressive MS cases and controls, with the hypothesis that S100A9 acts as a mediator of inflammation and neurodegeneration. We found that S100A9 expression was increased in MS, primarily localised to intravascular monocytes and extracellular, perivascular amyloid-like deposits, with these patterns correlating with markers of BBB damage and neurodegeneration. Conversely, when found in microglia, S100A9 was associated with reduced fibrin(ogen) deposition, a marker for BBB damage, and increased neuronal density, while its toxic phosphorylated form was reduced. These findings highlight a dual and compartment-specific role of S100A9 in progressive MS, with opposing effects depending on its localisation.

Quantitative analyses confirmed that total S100A9 expression was increased in both NLGM and NLWM in MS compared to controls. Within the NLGM, S100A9 was particularly enriched in the infragranular cortical layers, areas previously shown to be more susceptible to inflammatory damage in MS due to their distinct vascular architecture and metabolic demands^33,34^. This enrichment aligns with our prior observations of increased fibrin(ogen) leakage relating to neuronal loss in these layers in progressive MS^2^, reinforcing the concept that vascular dysfunction is a driver of neurodegeneration in the cortex. Of note, although previous studies described the presence of S100A9 in active demyelinating lesions^19–21^, we found comparable S100A9 expression in lesional and non-lesional areas. This may reflect the inactive stage of cortical demyelination in progressive MS, where active inflammatory cells are scarce^35,36^. A similar pattern has been observed for other DAMPs such as small heat shock proteins (sHSPs), which are equally expressed in lesional and non-lesional grey matter^37^. These findings suggest that S100A9 contributes to cortical pathology independently of demyelination and may reflect chronic inflammation.

S100A9 was primarily localised to intravascular monocytes and perivascular areas, consistent with previous studies reporting S100A9 expression in peripheral immune cells in MS^20,21^, and its aberrant amyloid deposition in AD and Parkinson’s disease^38,39^. Importantly, this represents the first evidence of a potential toxic role of extracellular amyloid aggregates in MS pathology, where perivascular S100A9 expression was related to BBB damage, as evidenced by its relationship with fibrin(ogen) accumulation. Its vascular origin is supported by the observation that both S100A9 monocytes and perivascular S100A9 plaques relates to each other and with reduced brain weight, linking vascular S100A9 compartmentalisation to cortical atrophy. Of note, these findings align with previous work showing the role of S100A9 in vascular-related disorders including atherosclerosis^40^, stroke^41^, and vascular dementia^42^.

The amyloid-like aggregation of S100A9 around vessels raises the possibility that this protein not only marks but actively contributes to vascular damage and inflammation. In AD, S100A9 can self-assemble into amyloid fibrils and promote a neuroinflammatory cascade^11^, with similar mechanisms suggested in TBI^14^. In addition, herpesviruses such as Epstein–Barr virus (EBV), recently confirmed as a key trigger of MS^43,44^, are known to upregulate S100A9 expression in monocytes and B cells^45,46^, potentially linking peripheral viral activation to MS pathology. These data suggest that peripherally derived S100A9 may enter the central nervous system through a disrupted BBB and act as both a DAMP and amyloid-forming protein, perpetuating inflammation and tissue damage. This mechanism is highly relevant in MS and warrant further investigation, as chronic vascular damage and smouldering inflammation are hallmarks of progressive disease^47^.

In contrast, microglial S100A9 expression was associated with protective features, including neuronal preservation and reduced fibrin(ogen) deposition. While S100A9 has classically been linked to pro-inflammatory functions via TLR4 and RAGE^48,49^, emerging evidence suggests that its activity is highly context- and dose-dependent. Studies show that low levels of S100A9 can promote microglial phagocytosis, inhibit excessive inflammation and contribute to tissue repair^16,50^, whereas high concentrations exacerbate inflammatory responses, as exemplified in age-related macular degeneration^51^. These dose-dependent effects may reflect different intracellular signalling pathways, post-translational modifications, and interactions with other inflammatory mediators. This is supported by the specific low expression of phosphorylated S100A9 (pS100A9) we observed in microglia from MS cases. Indeed, phosphorylation of S100A9, particularly at threonine 113, enhances its pro-inflammatory and amyloidogenic properties^52,53^. Therefore, the predominance of unphosphorylated S100A9 in microglia may represent a beneficial adaptation, preserving immune surveillance while limiting neurotoxicity in progressive MS. Consistent with this, we observed that TREM2+ microglia, a phenotype generally linked with anti-inflammatory and protective functions^54^, were increased in MS cases with high microglial S100A9 expression. This suggests that microglial S100A9, in its non-phosphorylated form, may align with a TREM2+ protective microglial program.

Importantly, we did not observe differences in microglial S100A9 expression between MS and controls, indicating that microglial S100A9 expression is likely an intrinsic component of the homeostatic microglial repertoire rather than a disease-induced marker. This aligns with previous studies listing S100A9 among homeostatic microglial genes, alongside TREM2, in non-diseased tissue^55,56^. What appears to differ across MS cases is whether microglia retain or lose this protective state. Cases maintaining this programme showed preserved neuronal density and minimal perivascular S100A9 plaques, whereas cases lacking it exhibited prominent perivascular S100A9 plaques, increased fibrin(ogen) accumulation, and reduced neuronal density. These findings suggest that a subset of MS cases may lose this intrinsic protective microglial pattern, potentially resulting in inadequate S100A9 clearance and increased extracellular aggregation. For instance, this is consistent with the established role of TREM2+ microglia in restricting amyloid accumulation in neurodegenerative models^57,58^. Whether microglia internalise vascularly derived S100A9 or instead retain microglial S100A9 intrinsically remains uncertain, and future investigation should clarify this distinction, essential to understanding the divergent microglial states that shape neurodegeneration in progressive MS.

We acknowledge that this study has limitations. The use of post-mortem tissue provides only a static view of complex and dynamic processes. Further, although dual roles of S100A9 have been described in other neurological conditions, this remains undefined in progressive MS. Finally, our study was restricted to the motor cortex, and it remains unclear whether similar compartmentalisation of S100A9 exists in other MS-affected regions such as other brain cortices, the hippocampus, or spinal cord. Despite these caveats, the robustness of our detailed neuropathological assessments, the consistent findings across multiple pathological readouts, and the alignment of our results with established mechanisms in other diseases strengthen the validity and relevance of our findings. Future studies should assess whether modulating S100A9 phosphorylation or blocking its extracellular aggregation can alter neurodegeneration or inflammation. Equally important will be exploring its effects on endothelial integrity, calcium homeostasis, and its potential interplay with viral triggers such as EBV. Longitudinal imaging and fluid biomarker studies, alongside experimental models, will be essential to determine whether targeting S100A9 could serve as a disease-modifying strategy in progressive MS.

In summary, this study sheds light on the multifaceted and context-specific roles of S100A9 in the progressive MS motor cortex. Our data support a model in which S100A9 acts as both a driver of neurodegeneration through vascular-associated amyloid-like aggregation, and as a potential neuroprotective agent within parenchymal microglia. Disentangling these roles may not only enhance our understanding of MS progression but may also inspire compartment-targeted therapeutic strategies that recalibrate inflammation, preserving beneficial microglial functions while neutralising harmful perivascular aggregation. Such approaches may help address the unmet therapeutic needs of progressive MS, where current treatments remain largely ineffective.

## Declarations

## Ethics approval & Acknowledgments

The authors acknowledge the UK MS Society tissue bank, and donors of human material, for their contribution to this work. All autopsy material was obtained with the relevant ethics committee approval (REC 08/MRE09/31+5) according to the regulations outlined in the UK Human Tissue Act (2004).

The authors thank Emeritus Professor Margaret M. Esiri for critical discussions.

## Consent for publication

## Data availability and statement

The data that support the findings of this study are available from the corresponding author upon reasonable request.

## Authors’ contributions

JP & GDL conceived and designed the study. JP, JA & RY acquired the data. JP, and MP performed statistical analyses. JP & GDL drafted the original manuscript, and all authors commented on the final manuscript..

