## Supplementary data for "The unexpected dual role of S100A9 amyloid protein on neurodegeneration in progressive multiple sclerosis motor cortex"

*Correspondence to:

Jonathan Pansieri, PhD

**Running Title : S100A9 expression in progressive multiple sclerosis motor cortex**

| Target | Primary Antibody | Antibody dilution | Target | Antigen Retrieval | Incubation Settings |
|---|---|---|---|---|---|
| PLP | Biorad #MCA839G | 1:1000 | Myelin | Citrate pH6 microwave | 1h RT |
| S100A9 | Abcam #ab92507 | 1:3000<br>1:1500(IF) | S100A9 | Citrate pH6 microwave | ON 4°C |
| PhosphoS100A9 (Thr113) | Life Technologies #PA5118785 | 1:500 | Phosphorylated S100A9 | Tris-EDTA pH9 Autoclave | ON 4°C |
| TREM2 | Abcam #ab318262 | 1:100 | Microglia | Tris-EDTA pH9 Autoclave | ON 4°C |
| NeuN | Millipore #MAB377 | 1:400<br>1:250(IF) | Neurons | Citrate pH6 Autoclave | 1h RT<br>ON 4°C (IF) |
| Iba1 | Abcam #ab15690 | 1:250 (IF) | Microglia | Citrate pH6 Microwave | ON 4°C |
| TMEM119 | Biolegend #853302 | 1:1000 (IF) | Microglia | Citrate pH6 Microwave | ON 4°C |
| GFAP | Dako #Z0334 | 1:2500(IF) | Astrocytes | Citrate pH6 microwave | 1h RT |
| CD163 | Bio-rad #MCA1853 | 1:3000 (IF) | Monocytes | Tris-EDTA pH9 Autoclave | ON 4°C |
| CD31/34 | Agilent/Biorad #M082301 #MCAP547 | 1:25(IF)<br>1:50(IF) | Endothelium | Citrate pH6 microwave | ON 4°C |
| Fibrin(ogen) | Abcam #ab58207 | 1:5000 | Fibrin(ogen) | Tris-EDTA pH9 Autoclave | 1h RT |

**Supplementary Table 1. Antibodies used in staining procedures presented in this article.** (IF = immuno-fluorescence).

| | Cortical layers (NLGM) | Total S100A9 accumulation (% of area coverage) | | | S100A9+ monocyte-associated vessel density (vessels/mm$^2$) | | | Perivascular S100A9 plaques (plaques/mm$^2$) | | | Microglial S100A9 (score) | | |
|---|---|---|---|---|---|---|---|---|---|---|---|---|---|
| | | MS | control | p-value | MS | control | p-value | MS | control | p-value | MS | control | p-value |
| supragranular layers | I | 0,53 ± 0,12 | 0,26 ± 0,09 | 0,38 | | | | | | | | | |
| | II | 0,51 ± 0,08 | 0,40 ± 0,08 | 0,51 | 5,62 ± 0,49 | 4,64 ± 1,35 | 0,47 | 1,24 ± 0,24 | 0,56 ± 0,34 | 0,14 | 0,17 ± 0,05 | 0,09 ± 0,06 | 0,96 |
| | III | 0,61 ± 0,08 | 0,38 ± 0,14 | 0,15 | | | | | | | | | |
| Infragranular layers | IV | 0,60 ± 0,07 | 0,31 ± 0,11 | 0,08 | | | | | | | | | |
| | V | 0,58 ± 0,07 | 0,22 ± 0,06 | 0,007 | 7,28 ± 0,48 | 4,0 ± 0,97 | 0,009 | 1,13 ± 0,18 | 0,19 ± 0,12 | 0,003 | 0,29 ± 0,06 | 0,22 ± 0,11 | 0,97 |
| | VI | 0,58 ± 0,08 | 0,18 ± 0,05 | 0,009 | | | | | | | | | |

**Supplementary Table 2. Motor cortical S100A9 expression in MS non lesional grey matter compared to controls.** Presence of S100A9 in the motor cortex was mainly observed in intravascular monocytes, extracellularly deposited around the vessels, and within microglia. Layer-by-layer analysis revealed significantly higher levels of S100A9 in MS cases, restricted to infragranular layers of the non-lesional grey-matter. (Data is presented as ± SEM. Significant differences are highlighted in blue. N/A = not applicable).

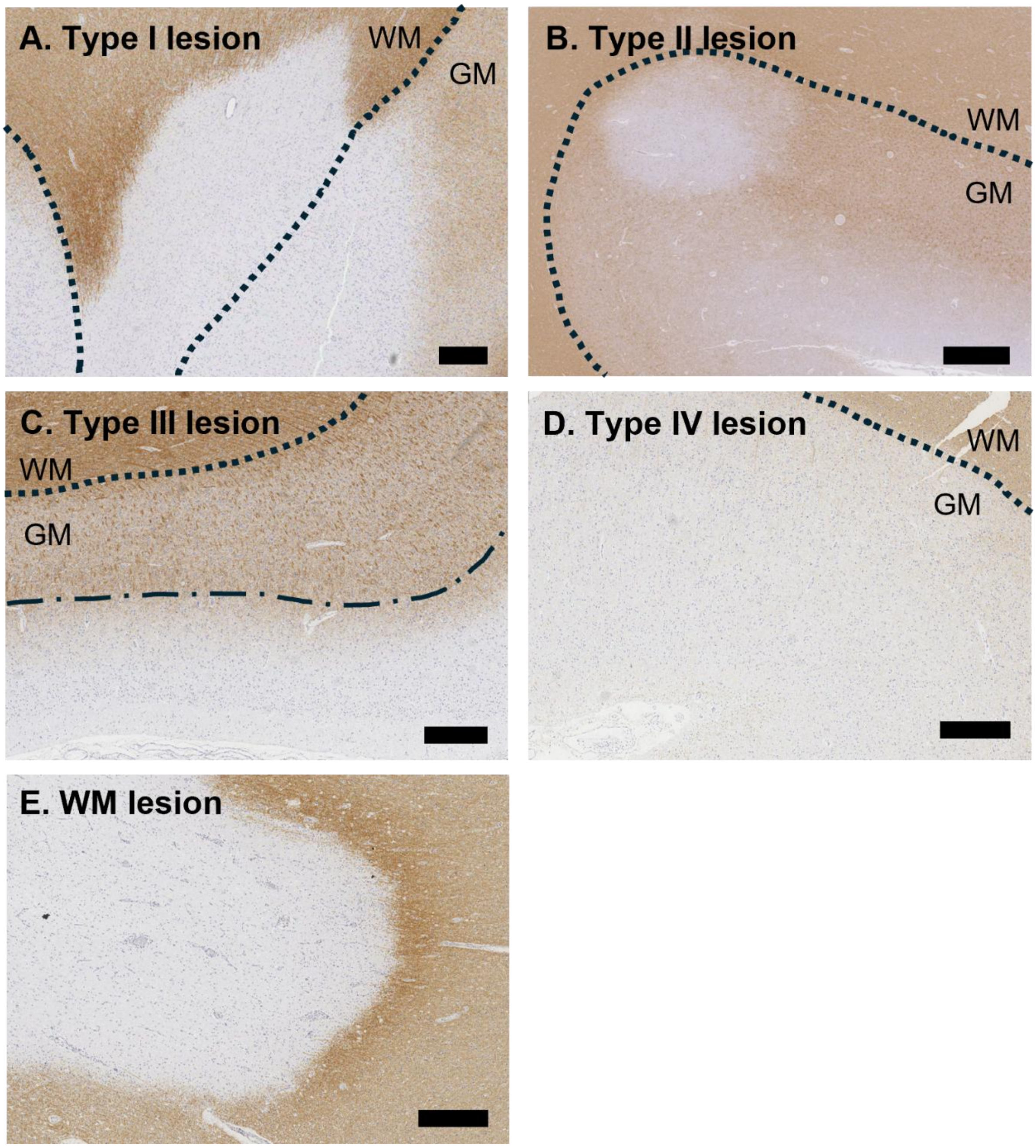


**Supplementary Figure 1. Demyelination in MS motor cortex.** Representative images of motor cortex stained with proteolipid protein (PLP) to identify total loss of myelin. **(A)** Type I demyelinated lesions (leukocortical) span both the white matter and the overlying cortical gray matter. **(B)** Type II demyelinated lesions (intracortical) are confined within the cortical gray matter of the motor cortex, without extending into the underlying white matter or to the subpial surface. **(C)** Type III demyelinated lesions (subpial) are found in the outermost layers of the cortical gray matter, directly beneath the pia mater, extending inward from the pial surface to supragranular layers, but do not penetrate infragranular layers. **(D)** Type IV demyelinated lesions (pan-cortical) are similar to type III lesions but do penetrate infragranular layers. **(E)** WM lesions are found in WM but do not penetrate GM. (Discontinued line in **(C)** represent supragranular/infragranular layers boundary ; dotted lines in **(A-D)** represent the WM/GM boundary; WM = white matter ; GM = grey matter ; scale bars are 500µm).

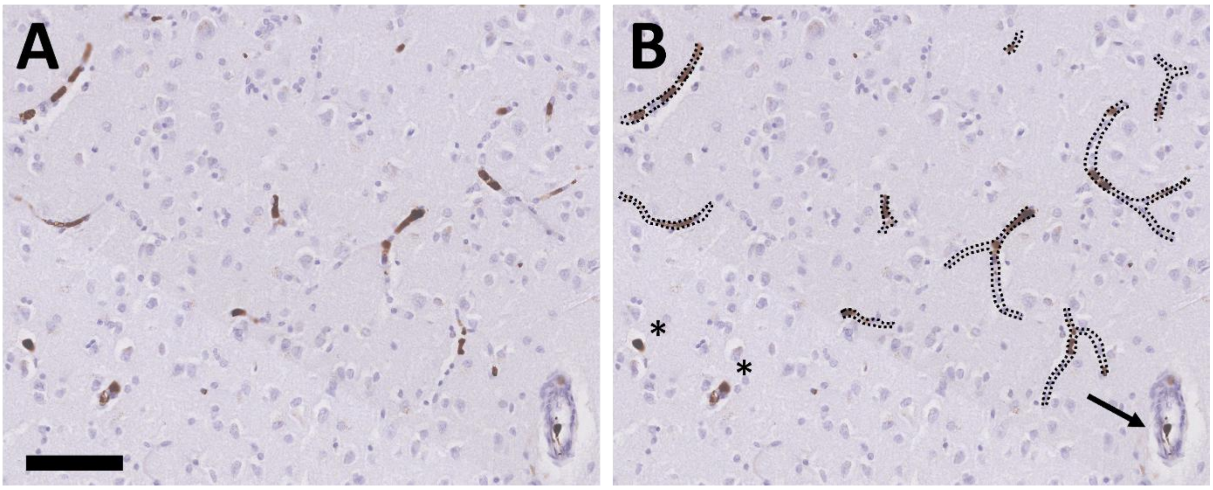


**Supplementary Figure 2. Assessment of S100A9+ monocyte-associated vessels in MS and control motor cortex.** **(A)** Vascular expression of S100A9 in a control case. **(B)** The presence of S100A9 in the grey matter of controls and MS non-lesional grey matter (NLGM) vessels was assessed in each field of view (FOV) by manually tallying the number of blood vessels exhibiting intravascular S100A9 chromogen-positivity and specific morphological characteristics (dotted lines). As illustrated, only vessels observed in the longitudinal plane were included in the count, revealing S100A9+ monocytes squeezed into elongated sausage-like shapes. Blood vessels in the transverse plane (asterisks) and structures as arterioles (arrow) were not counted. (Scale bar is 100 μm).

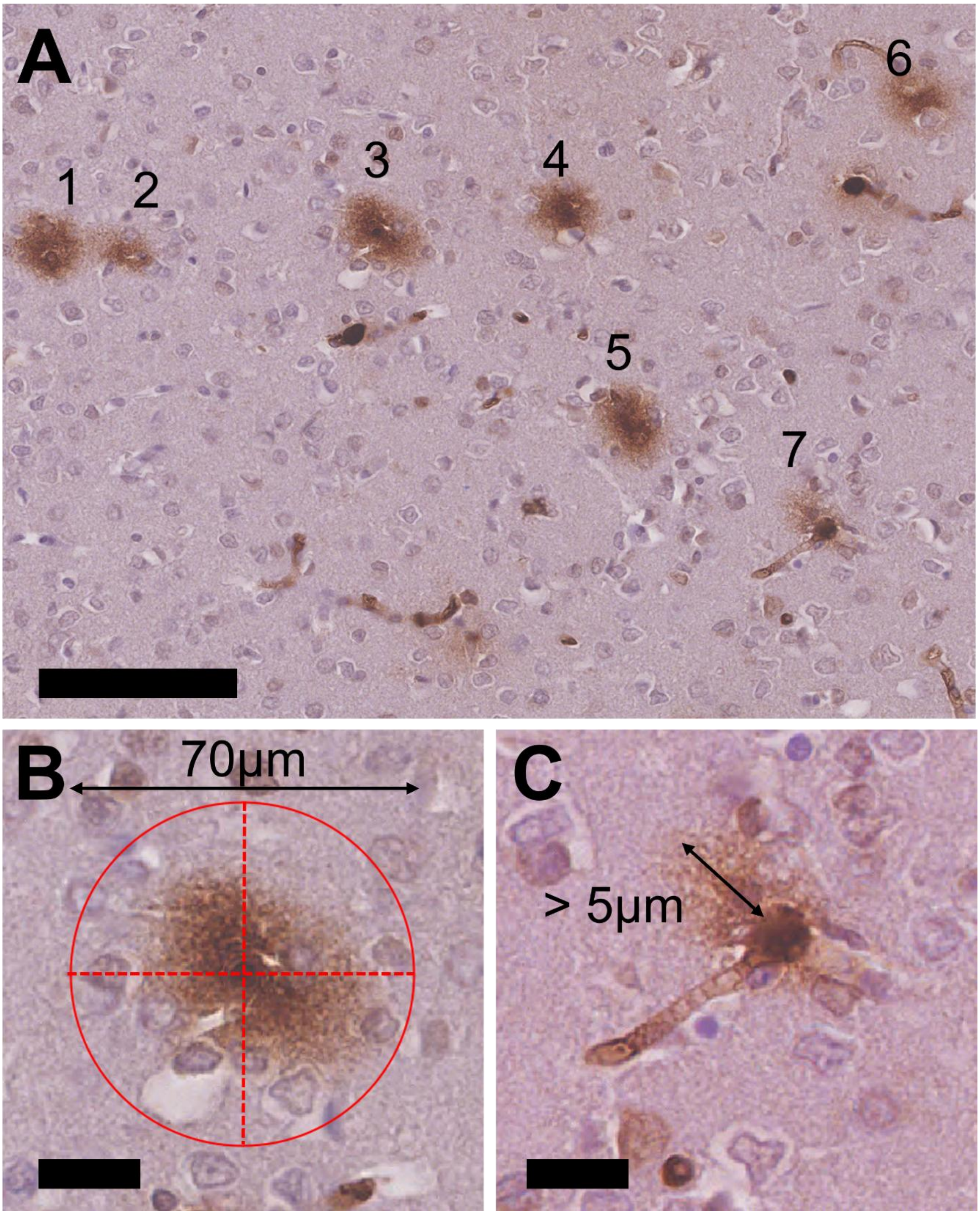


**Supplementary Figure 3. Assessment of perivascular S100A9+ plaques in MS and control motor cortex.** **(A)** Perivascular expression of S100A9 in the infragranular layer V of a MS case, where numbering illustrates the S100A9 plaques taken into account for quantitative analysis. The density of S100A9 plaques in the non-lesional grey matter (NLGM) **(B)** transverse and **(C)** longitudinal vessels was assessed in each field of view (FOV) by manually counting the number of plaques exhibiting S100A9 chromogen-positivity and expressed as plaques/mm$^2$. As illustrated, only chromogen-positive S100A9 plaques within **(B)** a 70µm radial distance from the vessel wall and showing **(C)** a diameter larger than 5µm were considered. (Scale bar in **(A)** is 100 µm, scale bars in **(B,C)** are 20 µm).

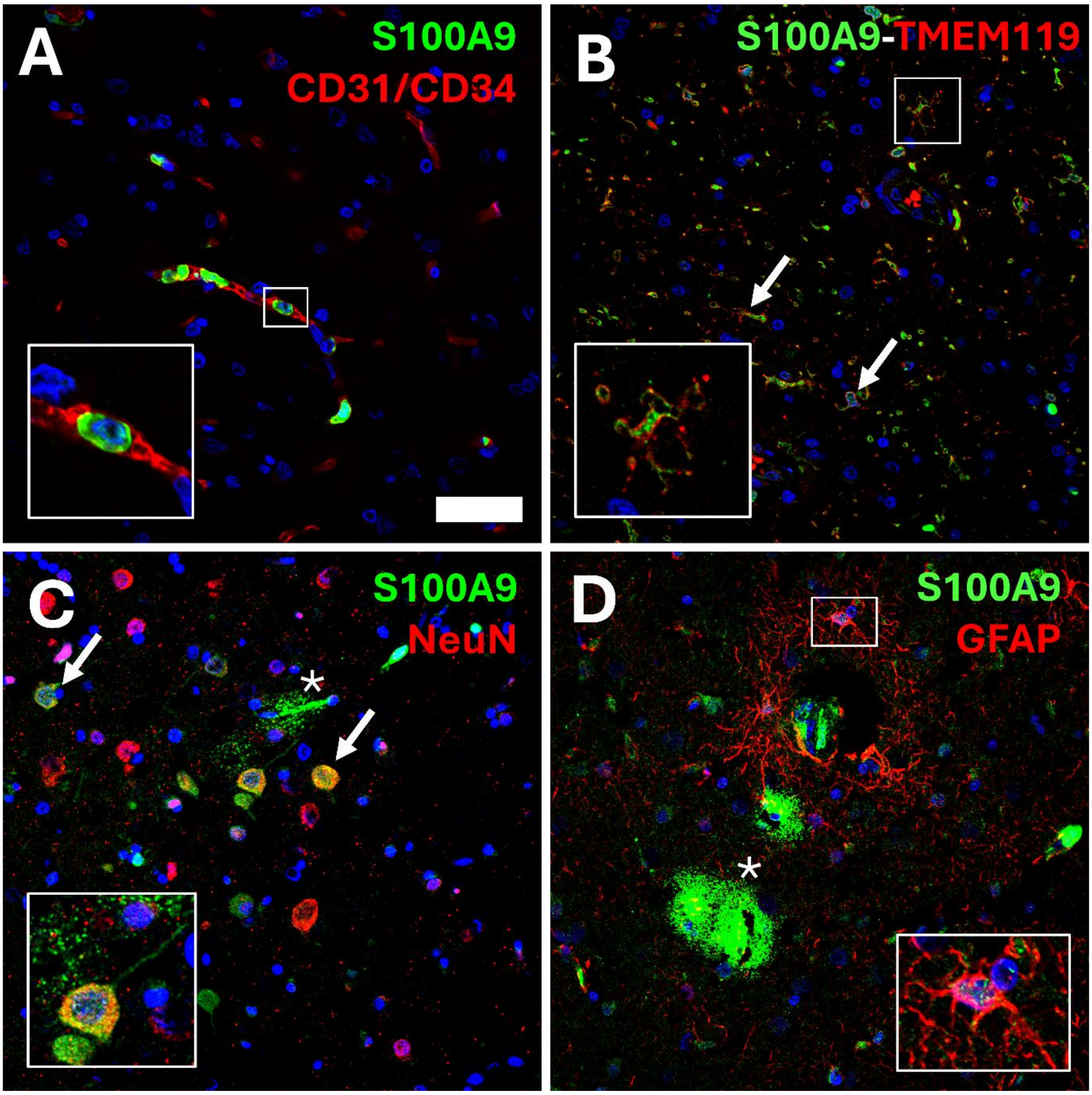


**Supplementary Figure 4. S100A9 is expressed in vessels and microglia, rarely in neurons, but not in astrocytes.** S100A9 (green) was compartmentalised in vessels (**A**, endothelial markers CD31-34, red) and expressed by TMEM119+ microglia (**B**, red), NeuN+ neurons (**C**, red) but rarely by astrocytes (**D,** GFAP+ astrocytes, **red**). Extracellular deposition can also be seen in Figure **D** (asterisks). (Scale bar is 50 µm, 400X magnification. Each picture is supplemented by Hoechst staining in blue which stain cell nuclei)

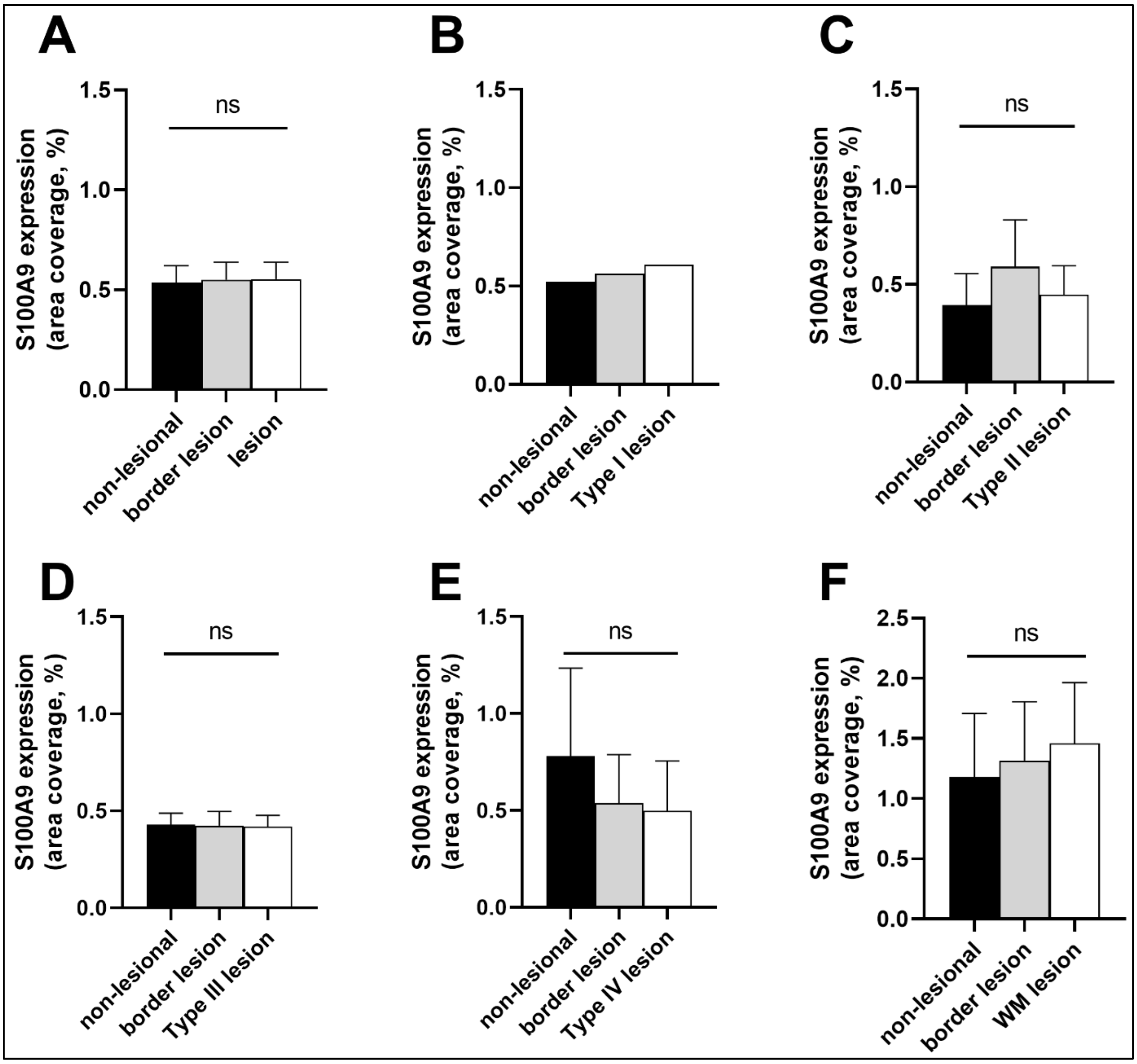


**Supplementary Figure 5. S100A9 expression and demyelination in MS cortex.** Comparison between non-lesional grey matter, non-lesional white matter and border lesional areas with **(A)** all demyelinated areas and **(B-F)** each type of demyelinated lesions shows no difference in S100A9 expression. Of important note, this analysis was stratified by comparing specific cortical layers and white matter distribution of the lesions with corresponding cortical layers or white matter non-lesional areas. (Friedman test with Dunn's correction for multiple comparisons ; ns= non-significant ; data presented as mean ± SEM, panel **B** represent only one lesion)

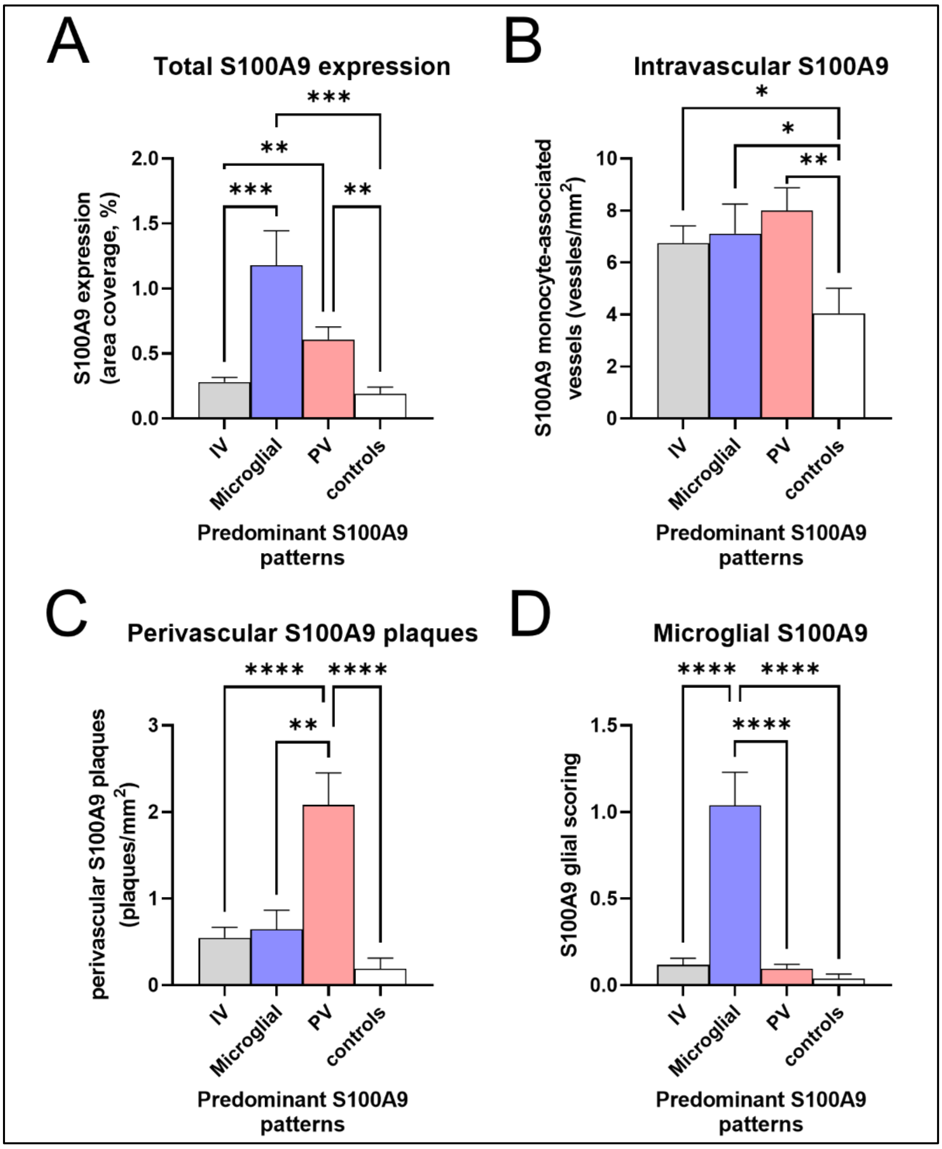


**Supplementary Figure 6. Quantitative and semi-quantitative measures of S100A9 depending on case-dependent patterns in MS cases, compared to control cohort. (A)** Increase in total S100A9 expression in MS compared to controls is driven by cases showing predominant microglial S100A9 and perivascular S100A9 plaques. **(B)** Increase in monocyte-associated S100A9 density in MS compared to controls is consistent for all case-dependent patterns. **(C)** Increase in perivascular S100A9 plaque density in MS compared to controls is driven by cases showing perivascular S100A9 expression. **(D)** Increase in microglial S100A9 expression in MS compared to controls is driven by cases showing predominant microglial S100A9 expression. (Results are presented as mean ± SEM ; *p<0.05; **p<0.01 ; ***p<0.001 **** p<0.0001 ; IV = intravascular S100A9 ; PV = perivascular S100A9 plaques ; NLGM = non-lesional grey matter, NLWM = non-lesional white matter)

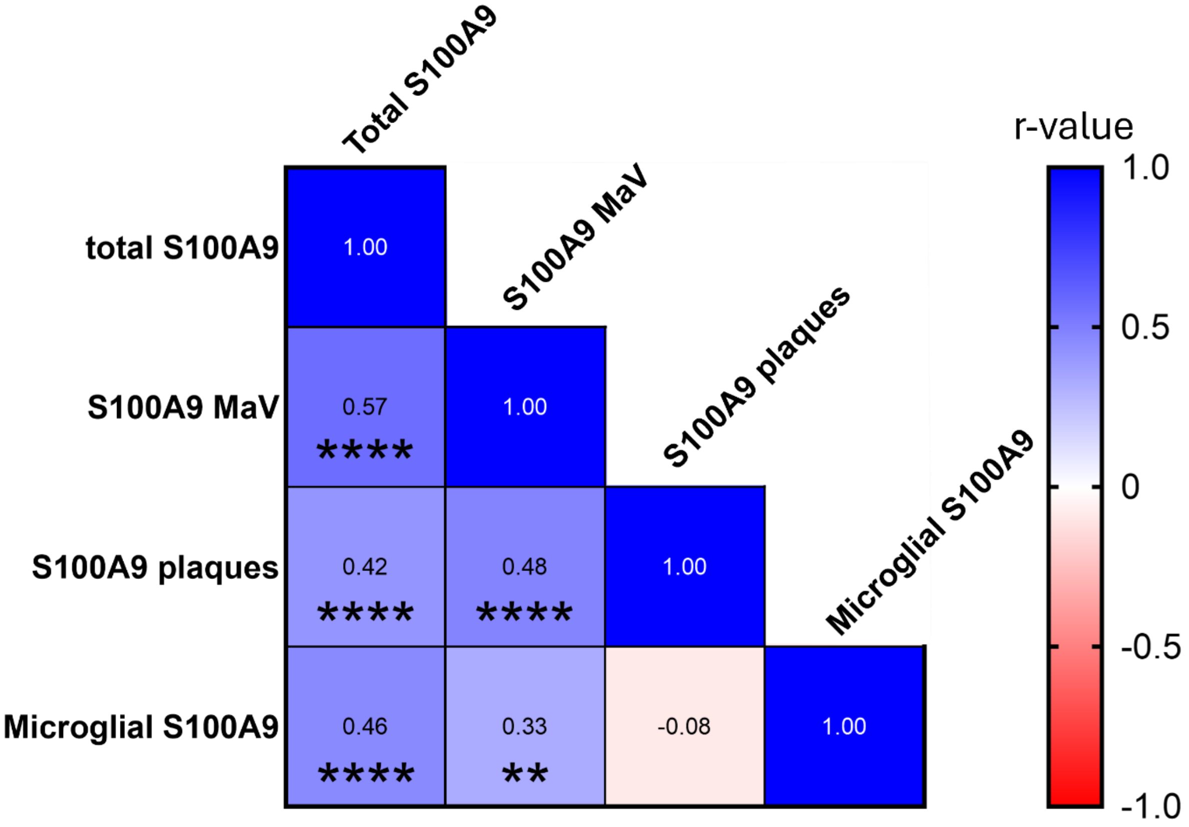


**Supplementary Figure 7. Spearman Correlation Matrix comparing quantitative and semi-quantitative measures of S100A9 expression.** A positive relationship between total S100A9 expression, S100A9+ monocyte-associated vessel density (r=0.57, p<0.0001), perivascular S100A9 plaque density (r=0.42, p<0.0001) and microglial S100A9 (r=0.46, p<0.0001) was found. In addition, S100A9+ monocyte-associated vessel density was associated with perivascular S100A9 plaque density (r=0.48, p<0.0001) and microglial S100A9 (r=0.33, p=0.007). Scale on the right represent the r-value scale, asterisks and data in each square illustrate the significance and the r value for each correlation, respectively. (Spearman correlation, **p < 0.01; **** p<0.0001 ; MaV = monocyte-associated vessel).

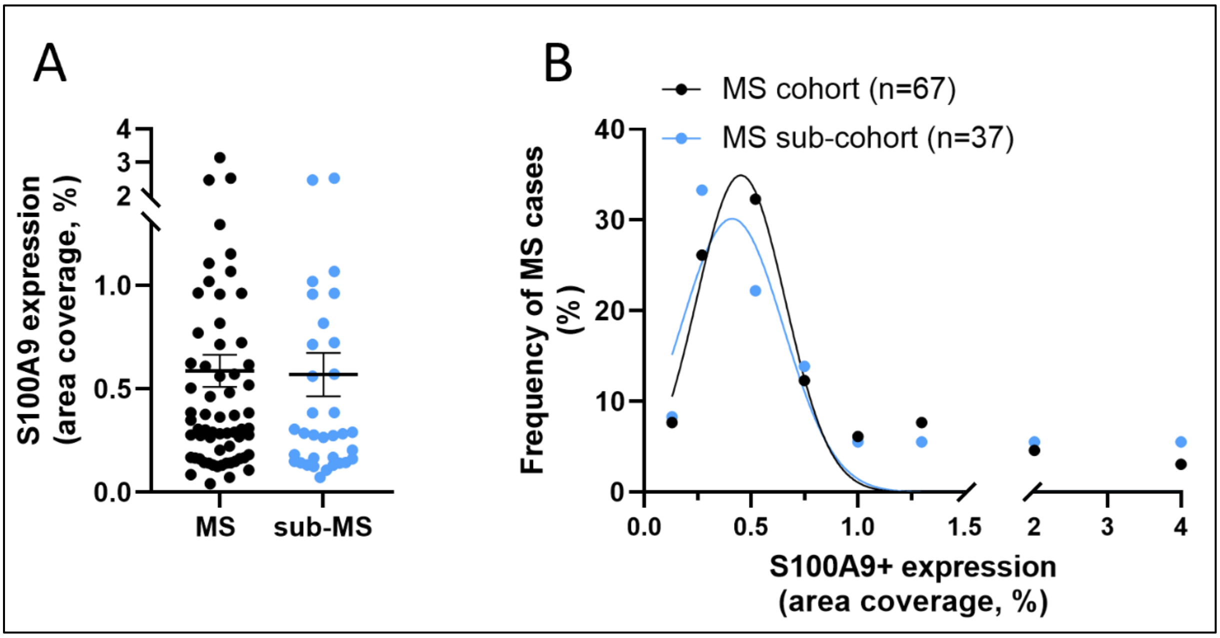


**Supplementary Figure 8. Case selection In MS cohort.** To investigate the relationships between S100A9 and brain cells, 37 MS cases which were previously well-characterised for various markers of interest were selected within the initial cohort of 67 MS cases. **(A)** No difference was found between the total MS cohort (n=67) and the subset (n=37, p >0.5), **(B)** following the same distribution in S100A9 expression over the cases.

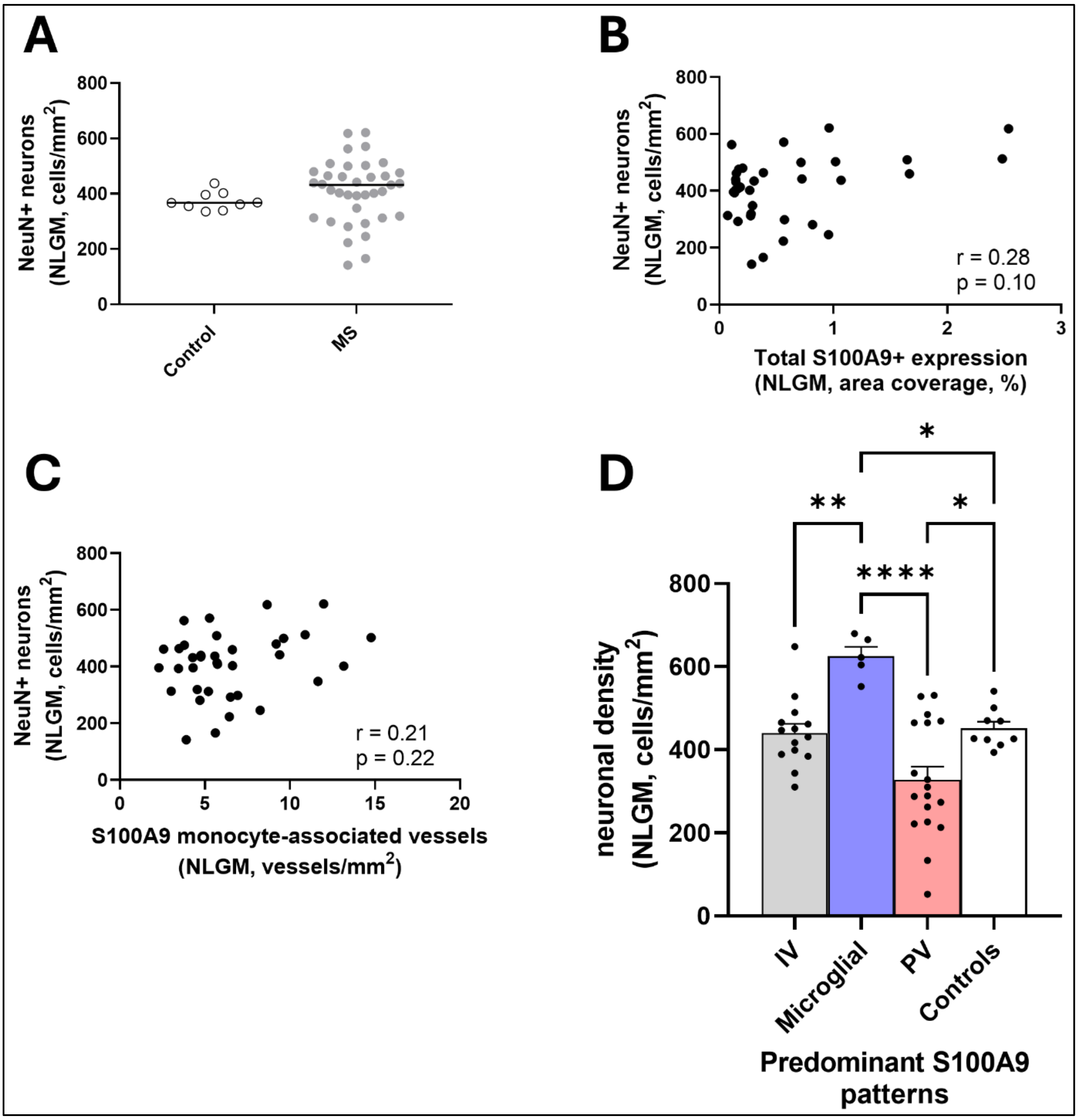


**Supplementary Figure 9. Neuronal densities in MS and controls. (A)** No difference in NeuN+ neuronal density was found in MS cases compared with controls. No relationship between NeuN+ neuronal density and **(B)** total S100A9 expression and **(C)** S100A9 monocyte-associated vessel density was found. **(D)** Comparing case-dependent S100A9 patterns in MS cases to controls, we found that cases with predominant microglial S100A9 show increased neuronal density, while cases with predominant perivascular S100A9 plaques show reduced neuronal density. (Results are presented as mean ± SEM ; *p<0.05; **p<0.01 ; **** p<0.0001 ; IV = intravascular S100A9 ; PV = perivascular S100A9 plaques ; NLGM = non-lesional grey matter, NLWM = non-lesional white matter).

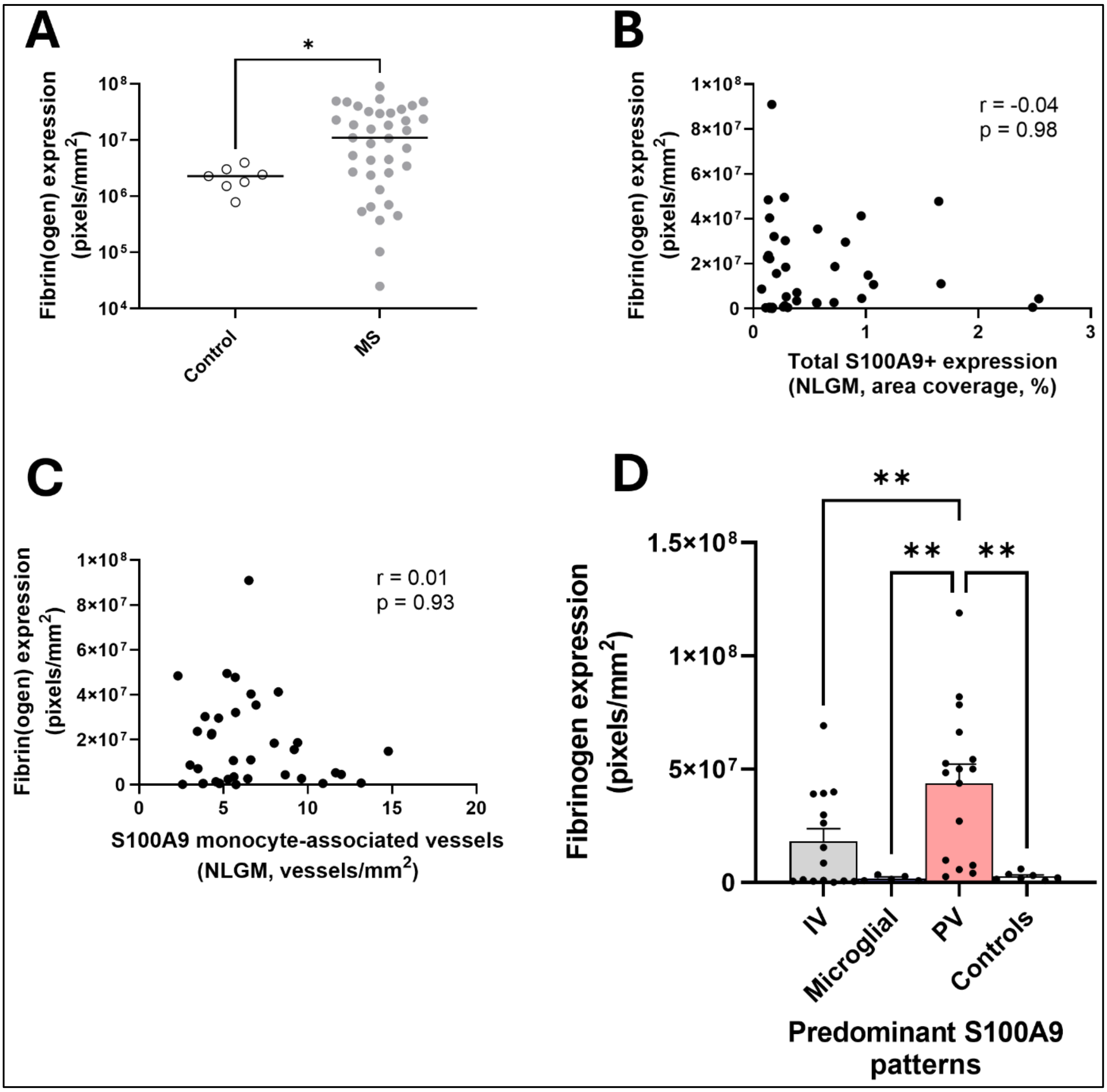


**Supplementary Figure 10. Fibrinogen expression in MS and controls. (A)** An increase in fibrin(ogen) expression was found in MS compared with control. No relationship between fibrinogen expression and **(B)** total S100A9 expression and **(C)** S100A9 monocyte-associated vessel density was found. **(D)** Comparing case-dependent S100A9 patterns in MS cases to controls, we found that cases with predominant microglial S100A9 show similar fibrinogen levels as controls, while cases with predominant perivascular S100A9 plaques show increased fibrinogen expression. (Results are presented as mean ± SEM ; *p<0.05; **p<0.01 ; IV = intravascular S100A9 ; PV = perivascular S100A9 plaques ; NLGM = non-lesional grey matter, NLWM = non-lesional white matter).

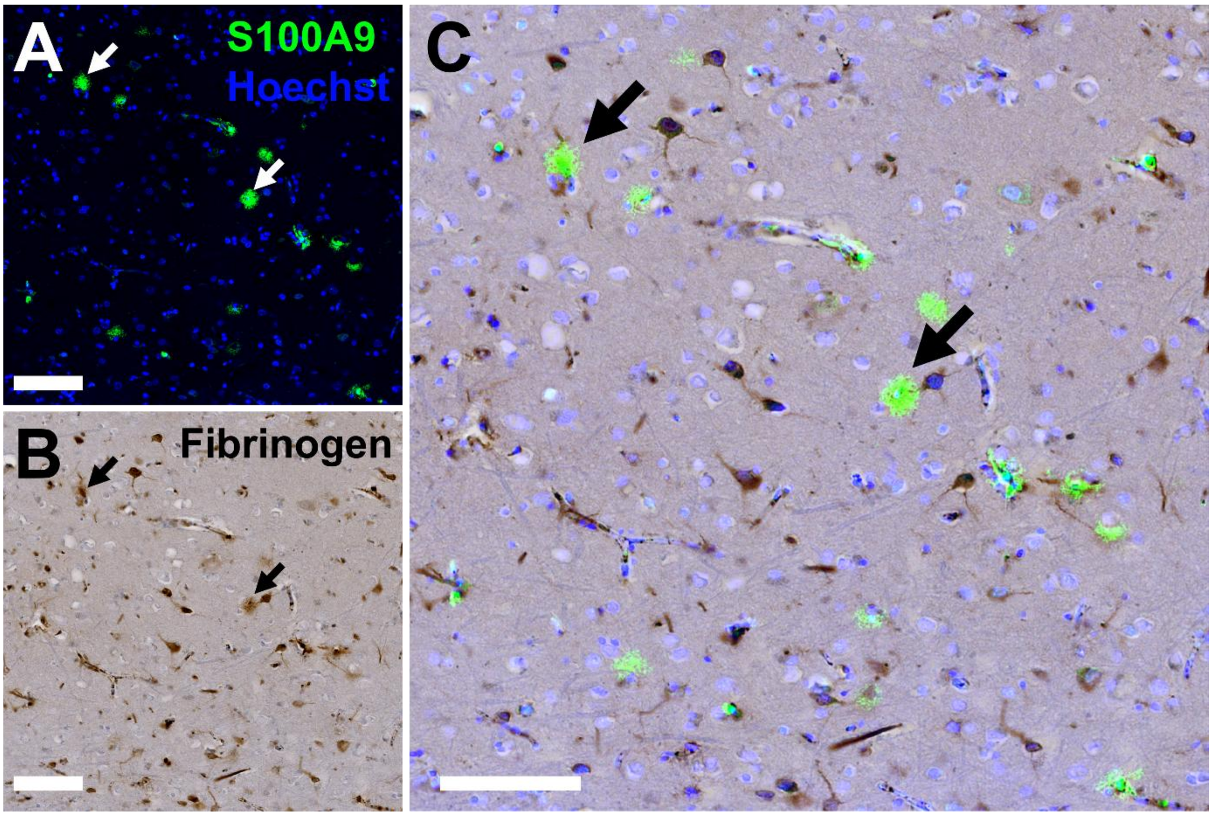


**Supplementary Figure 11. S100A9 colocalise with extracellular fibrinogen. (A)** S100A9 staining of MS motor cortex showing Perivascular S100A9 plaques (green). **(B)** Fibrinogen staining (brown) on the same section of MS motor cortex using DAB, showing extensive fibrinogen expression in various cell types and extracellularly deposited. **(C)** Combined pictures reveal co-location between extracellular fibrinogen and perivascular S100A9 plaques (arrows). (Scale bars are 100 µm, 400X magnification. Each fluorescent picture is supplemented by Hoechst staining in blue which stain cell nuclei)

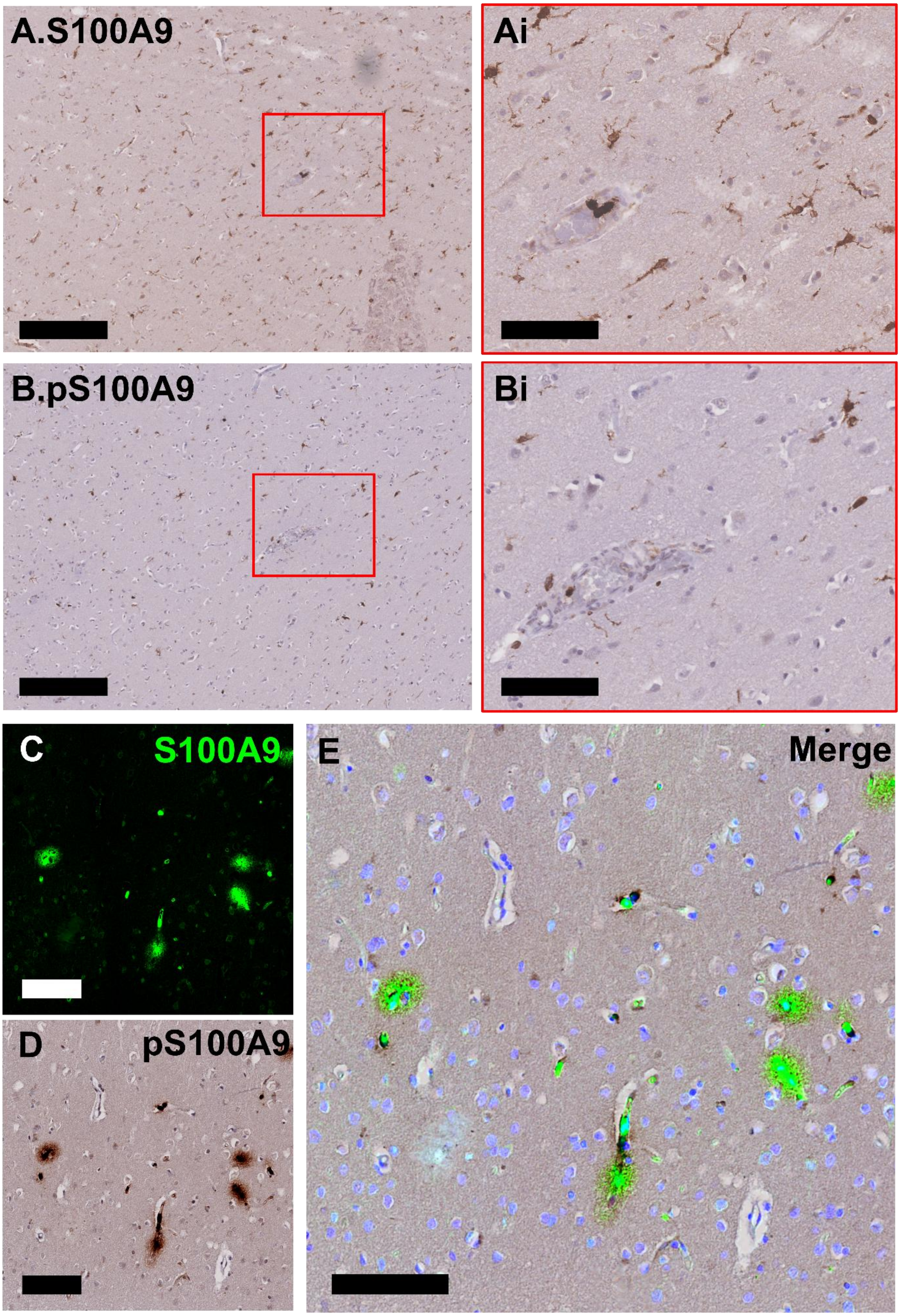


**Supplementary Figure 12. Phosphorylated S100A9 (pS100A9) is specifically reduced in microglial compartment. (A)** S100A9 is widely expressed in microglia of the MS motor cortex. Magnified picture in **(Ai)** illustrates both perivascular and parenchymal microglia expressing S100A9. Conversely, in the same FOV of an adjacent section from the same MS case, **(B)** pS100A9 expression in microglia is reduced and predominantly expressed in vessels and sparsely in perivascular microglia, as illustrated in magnified picture **(Bi)**. **(C-E)** Immunofluorescence/DAB co-labeling show similar S100A9 and pS100A9 expression in perivascular plaques. (Scale bars in **(A)** and **(B)** are 300 µm, scale bars in magnified pictures **Ai**, **Bi**, **C-E** are 100µm, fluorescent picture is supplemented by Hoechst staining in blue which stain cell nuclei).
